# Heterogeneity in effective size across the genome: effects on the Inverse Instantaneous Coalescence Rate (IICR) and implications for demographic inference under linked selection

**DOI:** 10.1101/2021.06.11.448122

**Authors:** Simon Boitard, Armando Arredondo, Camille Noûs, Lounès Chikhi, Olivier Mazet

## Abstract

The relative contribution of selection and neutrality in shaping species genetic diversity is one of the most central and controversial questions in evolutionary theory. Genomic data provide growing evidence that linked selection, i.e. the modification of genetic diversity at neutral sites through linkage with selected sites, might be pervasive over the genome. Several studies proposed that linked selection could be modelled as first approximation by a local reduction (e.g. purifying selection, selective sweeps) or increase (e.g. balancing selection) of effective population size (*N_e_*). At the genome-wide scale, this leads to variations of *N_e_* from one region to another, reflecting the heterogeneity of selective constraints and recombination rates between regions. We investigate here the consequences of such genomic variations of *N_e_* on the genome-wide distribution of coalescence times. The underlying motivation concerns the impact of linked selection on demographic inference, because the distribution of coalescence times is at the heart of several important demographic inference approaches. Using the concept of Inverse Instantaneous Coalescence Rate, we demonstrate that in a panmictic population, linked selection always results in a spurious apparent decrease of *N_e_* along time. Balancing selection has a particularly large effect, even when it concerns a very small part of the genome. We also study more general models including genuine population size changes, population structure or transient selection and find that the effect of linked selection can be significantly reduced by that of population structure. The models and conclusions presented here are also relevant to the study of other biological processes generating apparent variations of *N_e_* along the genome.

## Introduction

One of the greatest challenges of evolutionary biology is to understand how natural selection, mutation, recombination and genetic drift have shaped and are still shaping the patterns of genomic diversity of species living today (Charlesworth, 2010, Lewontin, 1974, Walsh and Lynch, 2018). In the last decade genomic data have become increasingly available for both model and non-model species. It is expected that by analysing these genomic data we will be able to better understand the respective roles of the different evolutionary forces (Charlesworth, 2010, Lewontin, 1974). In particular, it is believed that we will be able to identify the regions that have been shaped by selection, and those that may be more neutral (Johri et al., 2020, Pouyet et al., 2018). The relative importance of selection and neutrality in generating the genomic patterns of diversity we see today has been at the heart of many evolutionary debates and controversies over the last decades (Kimura, 1983, Lewontin, 1974, Ohta, 1992) and recent studies suggest that it still is (Comeron, 2017, Jensen et al., 2019, Kern and Hahn, 2018).

The concept of effective size (*N_e_*) is central to these debates (Charlesworth, 2009) because selection is expected to be more efficient when *N_e_* is large, and genetic drift to be the main driver of evolutionary change when *N_e_* is small (Ohta, 1992). For instance, Charlesworth (2009) notes that an autosomal locus under positive selection will behave neutrally when *s* < 1/4*N_e_*, where *s* is the selection intensity at this locus. At the same time it is commonly assumed that selection will itself imply a variation of *N_e_* across the genome (Charlesworth, 2009, Gossmann et al., 2011, Jiménez-Mena et al., 2016b). For instance, Gossmann et al. (2011) write that “*The effective population size is expected to vary across the genome as a consequence of genetic hitchhiking (Smith and Haigh, 1974) and background selection (Charlesworth et al., 1993)*”. They add that “*The action of both positive and negative natural selection, is expected to reduce the effective population size leading to lower levels of genetic diversity and reduced effectiveness of selection.*” They also stress that “*The evidence that there is variation in N_e_ within a genome comes from three sources. First, it has been shown that levels of neutral genetic diversity are correlated to rates of recombination in Drosophila […], humans […], and some plant species…*”. In his 2009 review on the concept of *N_e_* Charlesworth (2009) made a similar comment: “*N_e_ may also vary across different locations in the genome of a species […] because of the effects of selection at one site in the genome on the behaviour of variants at nearby sites*”. More recently, Jiménez-Mena et al. (2016a) stated that “*recent studies […] suggest that different segments of the genome might undergo different rates of genetic drift, potentially* ***challenging the idea that a single*** *N_e_* ***can account for the evolution of the genome***” (emphasis ours).

Under these explicit or implicit modelling frameworks, genomic regions with limited genetic diversity are thus seen as regions of low *N_e_* as a result of selective sweeps (Smith and Haigh, 1974) or background selection (Charlesworth et al., 1993), whereas regions with very high levels of genetic diversity may be seen as regions of large *N_e_* and could be explained by balancing selection (Charlesworth, 2009) (see also Hill and Robertson (1966)). Following that rationale, Jiménez-Mena et al. (2016b) suggested that different species might thus differ in the statistical distribution of *N_e_* across the genome and they presented such distributions for eleven species.

Given the central role played by the *N_e_* concept to detect, identify, and even *conceptualize* selection, it may be important, perhaps even enlightening, to explore the consequences of the ideas presented above with the concept of IICR (inverse instantaneous coalescence rate) recently introduced by Mazet et al. (2016). Indeed, the IICR is equivalent to the past temporal trajectory of *N_e_*, previously defined as the coalescent *N_e_* (Sjödin et al., 2005), in a panmictic population under neutrality, and it is the quantity estimated by the popular PSMC method of Li and Durbin (2011). The IICR was first defined by Mazet et al. (2016) for a sample size of two and its properties were studied under several models of population structure (Chikhi et al., 2018, Grusea et al., 2018, Rodríguez et al., 2018). It can also be used for demographic inference under neutrality and models of population structure (Arredondo et al., 2021, Chikhi et al., 2018). These studies showed that the IICR will significantly change over time when populations are structured, even when population size is actually constant. They also outlined that the IICR not only depends on the model of population structure but also on the sampling scheme, which questions the notion that an *N_e_* can be easily associated to (or is a property of) the model of interest when the model is structured (Chikhi et al., 2018, Rodríguez et al., 2018). The reason for this dependency is that the IICR is by definition a function of the distribution of coalescence times for two genes (*T*_2_), which is itself a function of both the evolutionary model and the location (in time and space) of the sampled genes.

One important assumption of the IICR studies mentioned above is that this distribution of *T*_2_ is homogeneous along the genome. The IICR, as defined and computed in previous studies, is thus a genomic average assuming that all loci follow a single Wright-Fisher model, with or without population structure, but with the same number of haploid genes. Whichever definition of *N_e_* one assumes, the underlying model assumes that *N_e_* is constant along the genome. If we now assume that *N_e_* varies across the genome as a consequence of selection (even as an approximation) then the variance of coalescence times should be different from that expected under a standard Wright-Fisher model, and the IICR should be a function of the underlying distribution of the *N_e_* values across the sampled genes. Genomic regions under different selection regimes might then exhibit specific signatures leading to differing IICR curves for each region. Alternatively, these regions might not be easy to identify but they might still influence the average genomic IICR estimated from sequenced genomes. In the present study we thus wish to explore ideas related to drift, selection and patterns of genomic diversity by studying the consequences of this putative genomic variation of *N_e_* on the IICR.

We first study the IICR under panmixia and constant population size but assuming that *N_e_* varies across the genome as a result of recurrent selection, using hypothetical distributions of *N_e_* and distributions inferred from genomic data. We then generalise the model to integrate temporal population size variations, population structure or transient selection effects. Finally, we compare IICR predictions with PSMC estimations obtained from simulated data under a model including variations of *N_e_* along the genome. Altogether, we advocate the use of the IICR as a concept that may help clarify what *N_e_* means and as one way, among others, to improve our understanding of the recent and ancient evolutionary history of species.

## The IICR under panmixia with several classes of (constant size) *N_e_* along the genome

### Methods: model description

We assume that the genome can be divided in *K* distinct classes, each of them characterized by a different *N_e_* that is constant over time. To model these differences of *N_e_*, we consider that each class *i* (*i* = 1 … *K*) evolves under a constant size Wright-Fisher (WF) model (i.e. panmictic with non-overlapping generations) with diploid population size *λ_i_N* (2 *λ_i_N* haploids), for some reference population size *N* corresponding to the actual number of diploids. Note that 2*N* represents an actual number of haploid genomes and that under the WF model, there is no ambiguity and *N* represents the *N_e_* under neutrality. Thus, *λ_i_* reflects the ratio of effective population size *N_e_* in class *i* relative to *N* and for convenience we may sometimes refer to *λ_i_* as *the* effective population size in class *i*. Assuming that *N* is large (i.e. that all *λ_i_N* are large), we rescale time by units of 2*N* generations and study the pairwise coalescence time resulting from this model. For two sequences sampled in the present (at time *t* = 0) for a locus from the *i^th^* class of the genome, we know from standard coalescent theory that the coalescence time 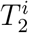 follows an exponential distribution with parameter 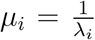, whose probability density function (pdf) is

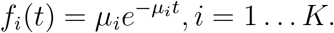

Denoting by *a_i_* the proportion of the genome corresponding to class *i*, the pdf of the coalescence time *T*_2_ at a random locus is thus

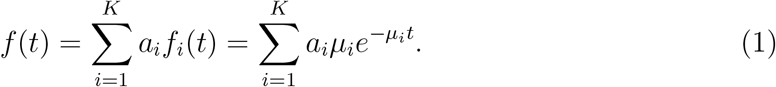

One may also see this distribution as the one we would obtain if we were able to sample a large number of independent coalescence times along the genome while covering each class *i* according to its true proportion *a_i_*. In the next section we study the properties of the IICR under this model.

### Results: IICR expression and main properties under panmixia

The IICR is a theoretical function that is intrinsically related to the expected distribution of coalescence times. Denoting *F* the cumulative distribution function of *T*_2_ for a given evolutionary model and sampling scheme, and *f* (*t*) = *F*′(*t*) its pdf, the IICR of a sample of size 2 is defined Mazet et al. (2016) as:

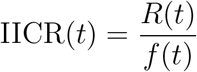

where

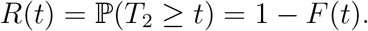

This theoretical quantity can be evaluated for any coalescent model by simulating a large number of independent *T*_2_ values and computing their empirical distribution (Chikhi et al., 2018). For a large class of models, it can also be obtained exactly using analytical or numerical approaches (Rodríguez et al., 2018). When analyzing a pair of real sequences, the evolutionary model that generated these sequences is unknown but the associated IICR can be estimated by SMC approaches like PSMC or MSMC (Schiffels and Durbin, 2013), which exploit the correlation structure of polymorphic sites along the genome to infer local coalescence times and their genome-wide distribution.

For our model with *K* different *λ_i_*, we have from equation (1):

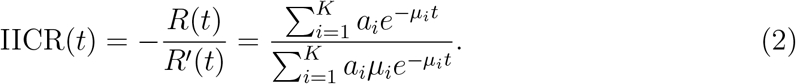

It is straightforward to see that the IICR is not constant as soon as there are at least two different values of *λ_i_* with non null proportion *a_i_* across the genome. To be more specific, we prove in the Supplementary Material that the IICR defined in formula (2) is *always increasing* from *t* = 0 to *t* = +∞ (i.e. backward in time). Thus, in a stationary panmictic population, the existence of at least two distinct *N_e_* across the genome (*λ_i_, i* > 1) is sufficient to infer a decreasing IICR (forward in time). In this situation, classical interpretations of PSMC plots under panmixia will lead to the wrong conclusion that the population size decreased through time. Alternatively, this signal could be (also wrongly) interpreted as the presence of population structure, since population structure can generate similar changes in the IICR (Mazet et al., 2016).

The magnitude of the IICR decrease can also be deduced from formula (2). Indeed, the value of the IICR at present is

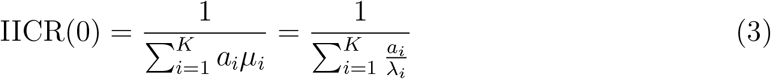

and the limit value when *t* → +∞ is equal to

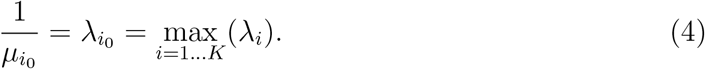

The present time value IICR(0) is thus necessarily between the smallest and largest *λ_i_*, as it is the harmonic mean of the *λ_i_*s weighted by their respective proportions *a_i_*. The asymptotic value IICR(+∞) is always the largest *λ_i_* found in the genome, *independent* of its proportion. In other words, even if a minute proportion of the genome has a high *λ_i_* due to balancing selection, under panmixia the IICR will necessarily plateau to this value in the ancient past. One intuitive explanation for the IICR growing (backward in time) towards the largest *λ_i_* is that the genes that are characterized by a large *N_e_* have much larger coalescence times than the rest of the genome. They thus contribute proportionately more to the most ancient part of the IICR curve.

### Results: a two-class panmictic model

These properties can be observed in Figure 1 where we represent the simplest case with *K* = 2 classes of genomic regions. In this figure we present the IICRs for *λ*_1_ = 0.1 and *λ*_2_ = 1, for proportions of *λ*_2_ (represented by the parameter *a*_2_) varying from 0 to 1. Consistent with the choice made in most studies inferring past population size changes, time is plotted in log10 scale in this Figure and all others shown in the main text.

**Figure 1:**
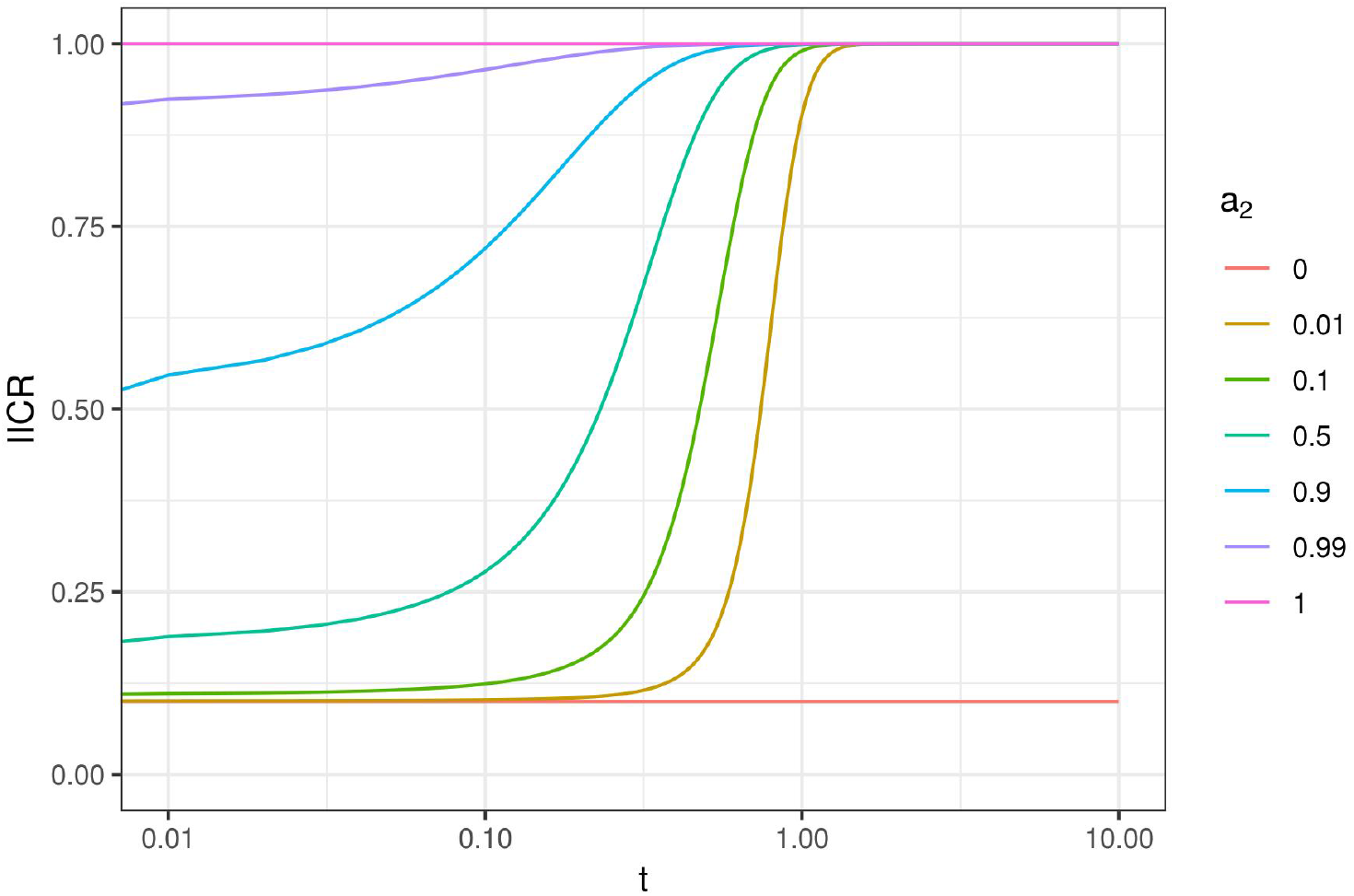
IICR curves for a panmictic model with *K* = 2 classes of genomic regions with constant size. Genomic regions of class *i* (*i* = 1, 2) have a constant population size *λ_i_N*, with *λ*_1_ = 0.1 and *λ*_2_ = 1. Their frequencies are *a*_1_ and *a*_2_, respectively, with *a*_1_ + *a*_2_ = 1. The IICR curves are represented for *a*_2_ values (representing neutrality, see main text) varying between zero and one. Time is plotted in log10 scale.

To simplify the interpretation of our results, we consider (by convention) throughout this manuscript that *λ_i_* = 1 corresponds to the neutral regions of the genome, whether *a_i_*, their relative proportion in the genome, is large or not. We thus do not necessarily consider that most of the genome is neutral in that sense. In this setting and in Figure 1, where *λ*_1_ = 0.1 and *λ*_2_ = 1, *a*_1_ can be interpreted as the fraction of the genome showing reduced *N_e_* by a multiplicative factor *λ*_1_ = 0.1 as a consequence of positive or background selection.

Figure 1 shows that for small values of *a*_2_ (i.e. when most of the genome is under *N_e_*-reducing selection) the IICR is S-shaped, slowly increasing backward from *λ*_1_ = 0.1 in the recent past to a plateau at *λ*_2_ = 1 in the ancient past. For increasing *a*_2_ values the IICR curves are becoming flatter as their left-most section flattens upward. Consistent with the properties outlined in previous section, these curves start (in recent times) at increasing IICR values above *λ*_1_ = 0.1 when the value of *a*_2_ increases, but the curves always reach the same ancient plateau at *λ*_2_ = 1. However, and this is an important point, this plateau is reached earlier as *a*_2_ increases. When *a*_2_=1, only the plateau remains and the IICR is flat at *λ*_2_ = 1 and when *a*_2_ = 0, it is a flat at *λ*_1_ = 0.1. Thus, when there is only one *λ_i_* over the genome, the IICR is constant over time and equal to that value, as expected for a population with constant size *λ_i_N* (Li and Durbin, 2011, Mazet et al., 2016).

If we now assume that the only type of selection present in the genome increases the effective size by an order of magnitude, with *a*_1_ and *a*_2_ corresponding to *λ*_1_ = 1 and *λ*_2_ = 10, we obtain exactly the same figure with the only difference that it is rescaled (Figure S1). This figure now shows that even if most of the genome is neutral, tiny amounts of *N_e_* increasing selection strongly influence the IICR, as it always grows backward towards the plateau corresponding to the largest of the two *λ_i_* values.

Altogether Figures 1 and S1 suggest that there is a strong asymmetry between selection reducing (background and positive) or increasing (balancing) *N_e_* in the genome in the way they affect IICR shapes. Balancing selection generates an ancient and high plateau at the level of *λ*_2_, even for small proportions of *a*_2_ (Figure S1), whereas positive and background selection generate a recent and relatively more modest decrease of the IICR for small values of *a*_1_, even assuming, as in Figure 1, that these generate a ten-fold decrease in *N_e_* (Figure 1).

### Results: a three-class panmictic model

To further explore the influence of both types of selection (reducing and increasing *N_e_*), we considered a model with 3 classes such that *λ*_1_ < 1, *λ*_2_ = 1 and *λ*_3_ > 1 (Figure 2). In this Figure we set the three *λ_i_* as (*λ*_1_*, λ*_2_*, λ*_3_) = (0.1, 1, 3). As above, *λ*_1_ < 1 corresponds to genomic regions under positive or background selection, *λ*_2_ = 1 corresponds to the neutral part of the genome and *λ*_3_ = 3 to genomic regions under balancing selection. In the left panel, we considered a fixed small proportion of balancing selection (*a*_3_ = 0.01), and allowed the proportions of neutral and positive or background selection to vary (*a*_1_ varied from 0 to 0.8, and thus *a*_2_ from 0.99 to 0.19). In the right panel, we considered a fixed and large proportion of positive or background selection (*a*_1_ = 0.5) and varied the proportion of regions under balancing selection (*a*_3_ from 0 to 0.1), and thus the proportion of neutral regions too (*a*_2_ between 0.5 and 0.4).

**Figure 2:**
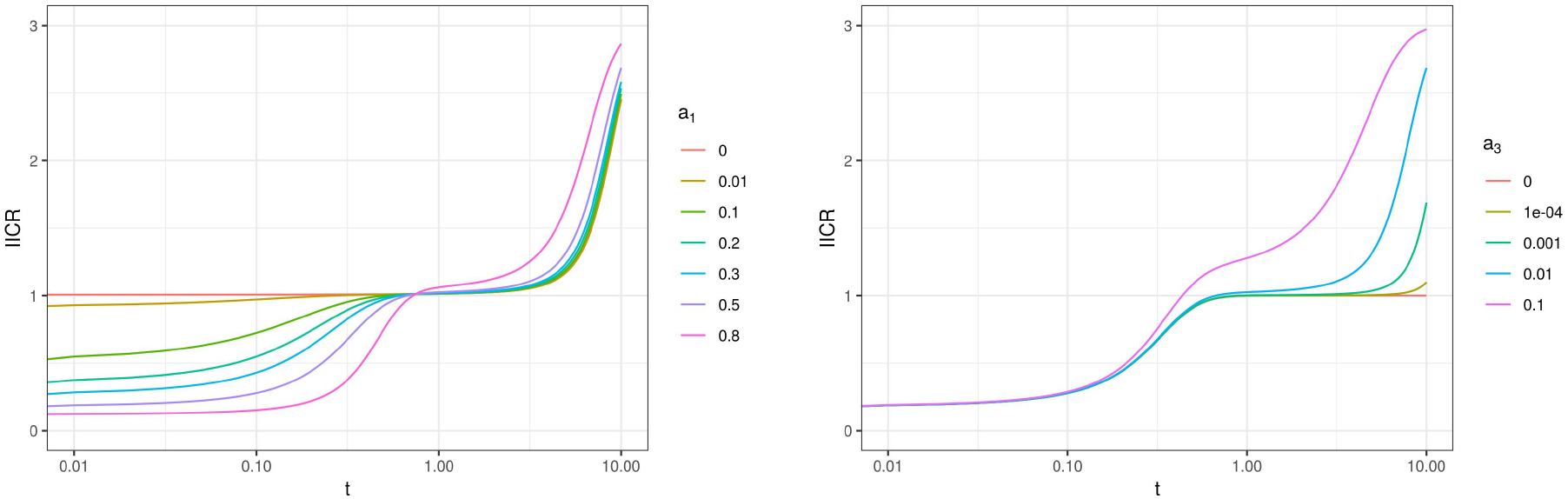
IICR for a panmictic model with *K* = 3 *λ_i_* values such that *λ*_1_ < 1, *λ*_2_ = 1 and *λ*_3_ > 1. The first class (or type) of genomic regions (*λ*_1_ < 1) is meant to represent regions of the genome under positive or negative selection and is modelled by a constant population size *λ*_1_*N* with *λ*_1_ = 0.1. Genomic regions of class 2 are meant to represent neutrality and they have a constant population size *λ*_2_*N* where *λ*_2_ = 1. Regions of class 3 are meant to represent genomic regions under balancing selection, they have a constant population size *λ*_3_*N* with *λ*_3_ = 3. Left panel: the frequency of class 3 is fixed at *a*_3_ = 0.01 and the frequencies of classes 1 and 2 are allowed to vary. The frequency *a*_1_ is given by the legend. Right panel: the frequency of class 1 is fixed at *a*_1_ = 0.5 and the frequency of classes 2 and 3 are allowed to vary. The frequency *a*_3_ is given by the legend.

Figure 2 shows similarities with Figure 1. Specifically, both figures suggest that regions reducing *N_e_* impact the IICR curves in the recent past whereas regions increasing *N_e_* impact the IICR in the ancient past. This is worth stressing given that our model assumes here that *N_e_* is reduced (in class 1) or increased (in class 3) in a stationary way throughout the genealogical history of the sampled genes (see the sections on transient selection for a different assumption). Also, small proportions of balancing selection seem to generate much bigger changes than small proportions of positive or background selection, as shown by the comparison of the IICRs obtained for *a*_1_ = 0.01 vs *a*_1_ = 0 on one hand (left panel) and for *a*_3_ = 0.01 vs *a*_3_ = 0 on the other hand (right panel).

There are however differences between Figure 2 and Figure 1. The simple fact that we consider both *N_e_*-reducing and *N_e_*-increasing forms of selection generates complex IICR curves, in which both forms of selection directly or indirectly impact the whole IICR curves. When neutral regions are frequent enough (*a*_1_ ≤ 0.5 and *a*_3_ ≤ 0.01), the IICR exhibits a plateau or a flattening at *λ*_2_ in its middle section, but for larger values of either *a*_1_ (left panel, *a*_1_ = 0.8) or *a*_3_ (right panel, *a*_3_ = 0.1) the proportion of neutral genomic regions decreases and the IICR curve only exhibits a short inflexion corresponding to *λ*_2_ = 1 before increasing backwards towards *λ*_3_. An interesting pattern related to this intermediate plateau is observed on the left panel when *a*_3_ is fixed: the IICR in the ancient past increases more and quicker (backward in time) for *a*_1_ = 0.8 than for lower values of *a*_1_, although *a*_1_ models the proportion of low *N_e_* regions in the region. This counterintuitive result likely comes from the fact that the proportion of neutral regions decreases when *a*_1_ increases, so that the IICR becomes more similar to that of a two class model with only *λ*_1_ and *λ*_3_, directly increasing to *λ*_3_.

Despite this complex interplay, Figure 2 provides some insights about our capacity to detect or quantify either type of selection based on the IICR. The left panel suggests that the IICR includes relevant information about the proportion of the genome under positive or background selection: for large values of *a*_1_, there is a quick decline of the IICR (forward in time) followed by a low plateau around *λ*_1_, whereas lower *a*_1_ values see a more recent and gradual decrease of the IICR without any clear recent plateau. However, this distinction is far less visible when plotting on a natural scale (Figure S2), in which case *a*_1_ values as different as 0.1 and 0.5 lead to quite similar IICRs. Besides, results on the importance of *a*_1_ are likely exaggerated by the small value of *λ*_1_ used in Figure 2, which implies a 10-fold reduction of *N_e_*. In comparison, our choice of *λ*_3_ only implies a 3-fold increase of *N_e_* in Figure 2.

While the value of *λ*_3_ (more generally of the highest *λ_i_*) determines the plateau of the IICR, the proportion of this class (*a*_3_) appears to determine to a large extent the speed of convergence (backward) to this ancient plateau (right panel). For the smallest *a*_3_ values (0.1 or 0.01%), this ancient plateau is not reached within the figure (for *t* ≤ 10) whereas a plateau corresponding to the neutral regions (*λ*_2_ = 1) is observed for quite long periods. For the largest *a*_3_ values considered here (1 or 10%), the convergence backward to the ancient plateau is so fast that the IICR does not exhibit the middle plateau around the neutral value, as already mentioned.

In any case, these results suggest that if selection can be seen as reducing or increasing *N_e_* in a panmictic population, the strongest effect on the IICR seems to be disproportionately the result of the largest *N_e_*, even though it may in practice affect ancient parts of the IICR curves that may not be easily reconstructed from real data. PSMC curves obtained from real data show a sharp decrease (forward in time) in the very ancient past in several species, including humans and Neanderthals. While this ancient decrease is usually ignored or interpreted as a statistical artefact resulting from the very low number of coalescence events dating back to this period, Figure 2 suggests that it is possibly due to divergent alleles maintained by balancing selection.

### Methods: distributions of *N_e_* inferred from real data

The above examples highlighted important and partly unexpected properties of the IICR when *N_e_* is variable along the genome. However, they relied on a very small number of classes with arbitrary *λ_i_* and *a_i_* values. It is thus not clear to which extent they inform us on the impact of linked selection in real species, where the combined variations of gene density, selection form or intensity and local recombination rate generate complex *N_e_* distributions. In this section we consider two model species for which variation in *N_e_* has been documented or estimated, the fruit fly *Drosophila melanogaster* and humans (Figure 3).

**Figure 3:**
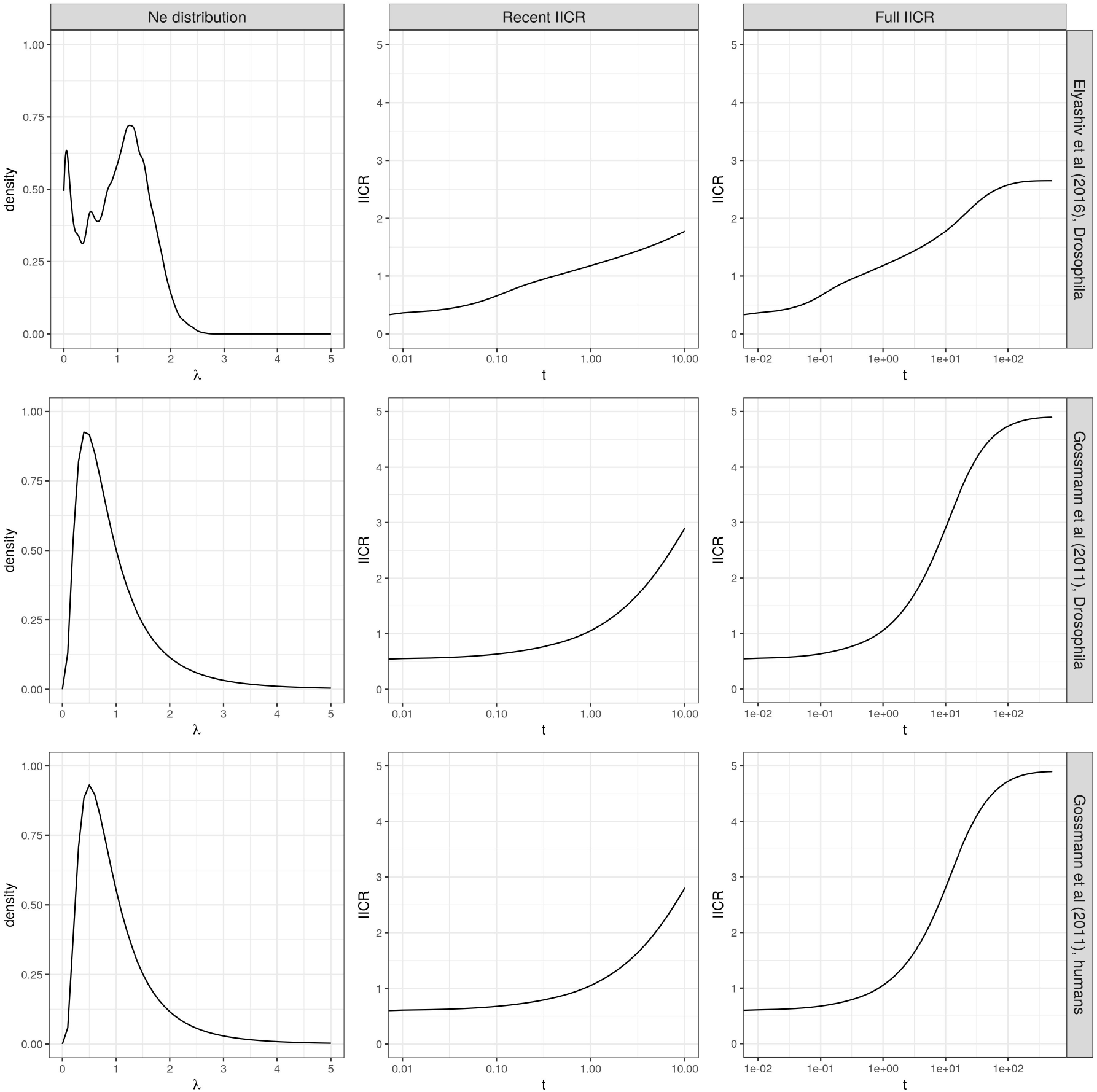
IICRs for panmictic models with large numbers of classes. This figure represents genome-wide distributions of *λ_i_* (left panels) and the associated IICRs until *t* = 10 (middle panels) or *t* = 500 (right panels). Top panels: IICR for *Drosophila melanogaster* (Raleigh, North Carolina population) based on the *N_e_* distribution estimated by Elyashiv et al. (2016). Middle panels: IICR for *D. melanogaster* (Zimbabwe population) based on the *N_e_* distribution estimated by Gossmann et al. (2011) assuming a lognormal distribution. To make the two IICRs comparable, the distribution estimated by Elyashiv et al. (2016) (top left) was re-scaled to have an average of one, as assumed in the analysis of Gossmann et al. (2011) (middle left). Bottom panels: IICR for humans (Yoruba population) based on the *N_e_* distribution estimated by Gossmann et al. (2011) assuming a lognormal distribution.

In the case of *Drosophila melanogaster*, we compared two different distributions of *λ_i_* over the genome, obtained by Gossmann et al. (2011) and Elyashiv et al. (2016). These two methods combine polymorphism data from the focal species and divergence data with closely related species, but they are based on very different approaches: the method of Elyashiv et al. (2016) explicitly models selection and its impact on the pairwise coalescence rate in each genomic region, while the method of Gossmann et al. (2011) assumes a log-normal distribution of *N_e_* over the genome and estimates its scale parameter from a large number of loci. For each of these two methods, the distribution obtained for *Drosophila melanogaster* was converted into a discrete distribution of *λ_i_* values with *K* = 25 and the associated IICR was computed using formula (2) (see the Supplementary Material for more details). As a comparison with another species, we also considered the distribution obtained by Gossmann et al. (2011) for humans.

### Results: distributions of *N_e_* inferred from real data

The distribution of *λ* inferred by Elyashiv et al. (2016) for *Drosophila* differed from the other two on two aspects (Figure 3). First, it had a lower support (up to *λ_i_* = 2.5, versus *λ_i_* = 5 for the others). This implied a smaller plateau of the IICR (as expected from equation (4)), but this effect was mainly visible at very ancient times (back to *t* = 500, right column) for which the IICR is unlikely to be observed from real data. Second, it had a mode for very low *λ_i_* values, which probably resulted from the inclusion of regions with very low recombination where the impact of linked selection is substantial. This mode had a limited effect on the IICR (see Figure S3 for an IICR obtained after filtering out *λ* values below 0.25 from the distribution).

Despite the differences between the species and the methods used to estimate the variation in *N_e_*, we obtained rather similar IICRs between *t* = 0 and *t* = 10 (middle column). The magnitude of the decrease observed in these IICRs was also comparable to that expected from Figure 2 for small values of *a*_1_ (e.g. *a*_1_ = 0.1, top right panel). Consequently, a long term 5 fold IICR decrease (from *t* = 10 to *t* = 0 forward in time) could realistically be the result, in both humans and *Drosophila melanogaster*, of a moderate proportion of loci with very small *N_e_* (Figure 2, *a*_1_ = 0.1, Figure 3, top) or from a larger proportion of loci with only slightly decreased *N_e_* (Figure 3, middle and bottom), all as a consequence of linked selection. Obviously, this conclusion can only be seen as a first order approximation, given that neither the estimation of the *N_e_* distribution by Elyashiv et al. (2016) or Gossmann et al. (2011), nor the computation of the resulting IICR, account for population demography or structure. Models including these aspects when computing the IICR are considered in the next section.

## Generalisation to more complex models

### Methods: extended model

We can generalise equation (2) to more complex models by still assuming that the genome is divided into *K* groups of loci each characterized by a different coalescence rate history. However, instead of describing this history by assuming panmixia and constant population size (*λ_i_*N), we can study different demographic models with departures from these assumptions, including models with panmixia and population size changes, models with population structure and models with transient (rather than recurrent) selection. In this more general framework, let us denote *f_i_*(*t*) the *pdf* of the coalescence time 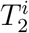. in the *i*-th class and *a_i_* the proportion of the genome in this class. The IICR is:

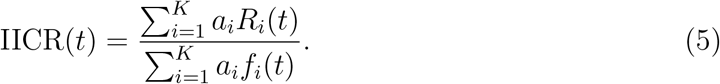

where 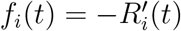.

### Results: panmixia and population size changes

One first potential application of this general framework is to study how linked selection interferes with genuine temporal variations of the population size. For instance, a natural question would be to know whether the spurious signal of recent population size decline arising from positive or background selection is strong enough to mask a genuine recent population expansion. To answer this question, we considered a simple extension of the two-class model studied in Figure 1 (*K* = 2, *λ*_1_ = 0.1 and *λ*_2_ = 1), where the population sizes in the two classes are multiplied by the same factor at a given time *T* before present. This expansion factor was set either to 5 in order to mimic the magnitude of (opposite) linked selection effects (Figure 4), or to 100 to mimic the very strong recent expansion that may be observed in some species including humans (Figure S4. The IICR of this model was computed by inserting known analytical expressions for the pdf of 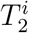 in each class *i* (e.g. (Mazet et al., 2015)) into formula (5). Note that the same approach could be applied to arbitrary complex demographic and selective scenarios, as long as the same temporal variations are applied to all classes.

**Figure 4:**
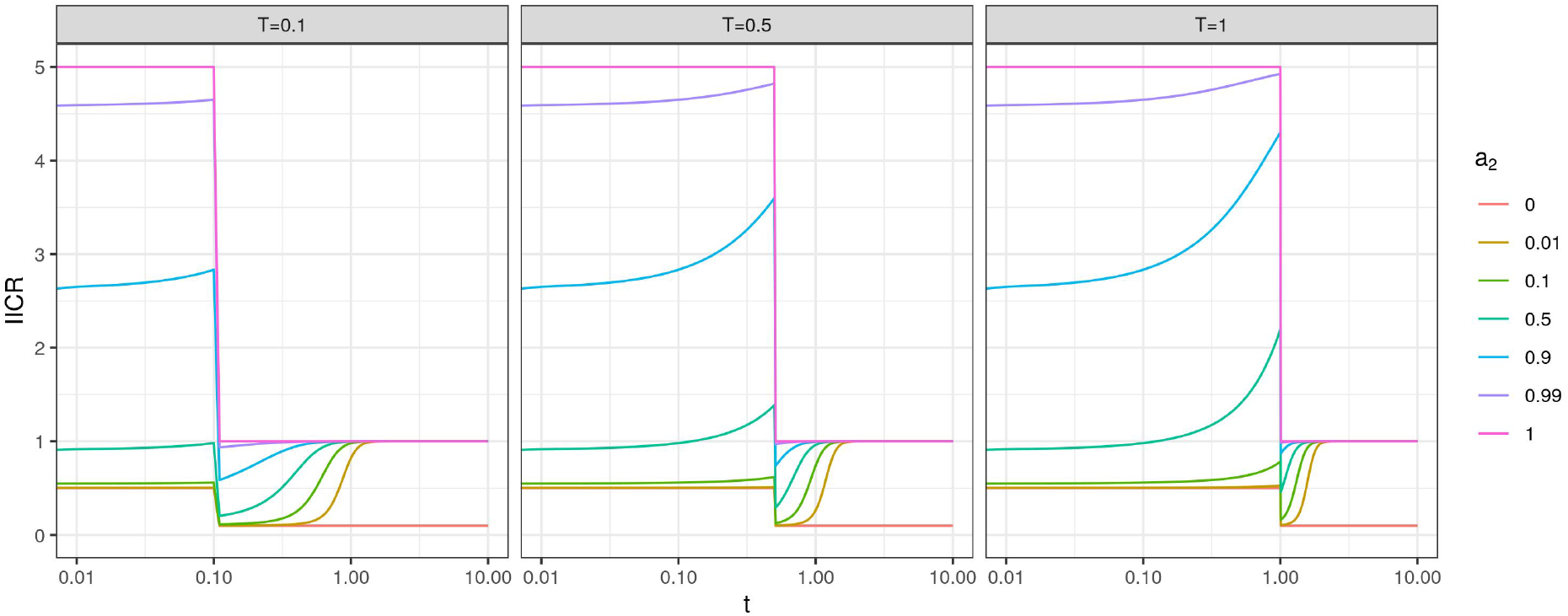
IICR curves for a panmictic model with a recent 5 fold expansion and *K* = 2 classes of genomic regions. Regions of class 1 and 2 have an ancestral population size 2*Nλ*_1_ and 2*Nλ*_2_ and a recent population size 10*Nλ*_1_ and 10*Nλ*_2_, with *λ*_1_ = 0.1 and *λ*_2_ = 1. Each panel corresponds to a different expansion time, indicated in the panel header. Frequencies *a*_1_ and *a*_2_ of the 2 classes are given by the legend (*a*_1_ + *a*_2_ = 1).

In the specific scenario considered here, we found that a strong proportion of selection in the genome could mask a genuine 5 fold expansion or even lead to the opposite conclusion of a population size decline (Figure 4). When 50% of the genome was under selection, the IICR showed transient temporal variations around the expansion time *T* (whose magnitude depended on *T*) but could at first approximation be interpreted as a constant population size history. When 90% of the genome was under selection, the overall pattern was that of a two fold decline. In contrast, smaller proportions of selection (10% of the genome or less) did not strongly affect the signal of population expansion. For stronger expansion events (100 fold, Figure S4), the IICR showed a significant increase for all values of *a*_1_ and *T*, but the IICR increase was much weaker than the true population size expansion: around 15 fold for *a*_1_ = 0.5 and 10 fold for *a*_1_ = 0.9. These results confirm that linked selection can significantly bias population size change inference, even in the presence of clear genuine demographic events.

### Results: stationary population structure

One other important extension of the models considered above is to account for population structure when modelling each genomic class. To illustrate this idea, we first considered a model with *K* = 2, *λ*_1_ = 0.1 and *λ*_2_ = 1 as in Figure 1. Here we assumed that these two classes evolved under a n-island model with the same number of demes (*n* = 10), the difference in *N_e_* being modelled through the use of different deme sizes in the two classes (*λ*_1_*N* and *λ*_2_*N*) We further assumed that selection did not affect migration, so that the *per* generation migration rate *m* was the same for the two classes. In other words, selection reducing *N_e_* is assumed to operate after migration and thus only affects coalescence rates, but not migration rates, of the two genomic regions. This implies that the scaled migration rate *M* = 2*Nm* is identical in the two classes (time scale is still 2*N* here, but *λ_i_N* now refers to deme diploid size rather than to the entire population size). One way of seeing this is by considering that there are 2*N* haploid genomes in each deme with scaled migration rate 2*Nm* and that selection acts on the different genomic regions by changing drift by a factor *λ_i_*.

As already mentioned and exploited in previous studies on the IICR (Grusea et al., 2018, Mazet et al., 2016, Rodríguez et al., 2018), the distribution of coalescence times under a symmetrical n-island model can be derived analytically (Herbots, 1994). Extending these derivations to a model with general deme size *λ_i_N*, instead of *N* in previous studies, we can show (see the Supplementary Material) that in this case

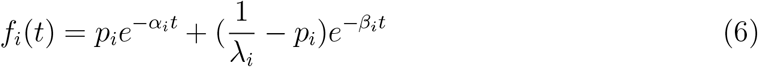

with

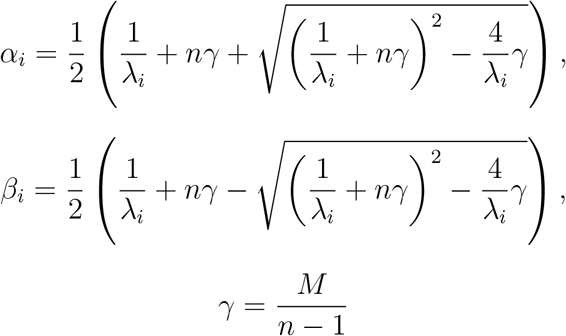

and

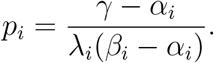

Setting *λ_i_* = 1 for all *i* recovers the results of Mazet et al. (2016). The IICR of an n-island model with two classes of deme size can be obtained by computing *f_i_*(*t*) with each *λ_i_* using Equation (6) and inserting the results into Equation (5).

IICR curves obtained for this two class n-island model are shown in Figure 5 for different values of the scaled migration rate. For *M* = 5, they are similar to those shown in Figure 1. This was expected given that an n-island model with high migration (*M* ≫ 1) should behave in a way that is similar to a panmictic model with population size *Nn*, except in the recent past where the IICR of the n-island still reflects local deme size (Mazet et al., 2016). For lower migration rates, the two extreme models with *a*_2_ = 0 (red curve) or *a*_2_ = 1 (violet) show that a higher plateau of the IICR is observed as *M* decreases, which was again expected (Mazet et al., 2016).

**Figure 5:**
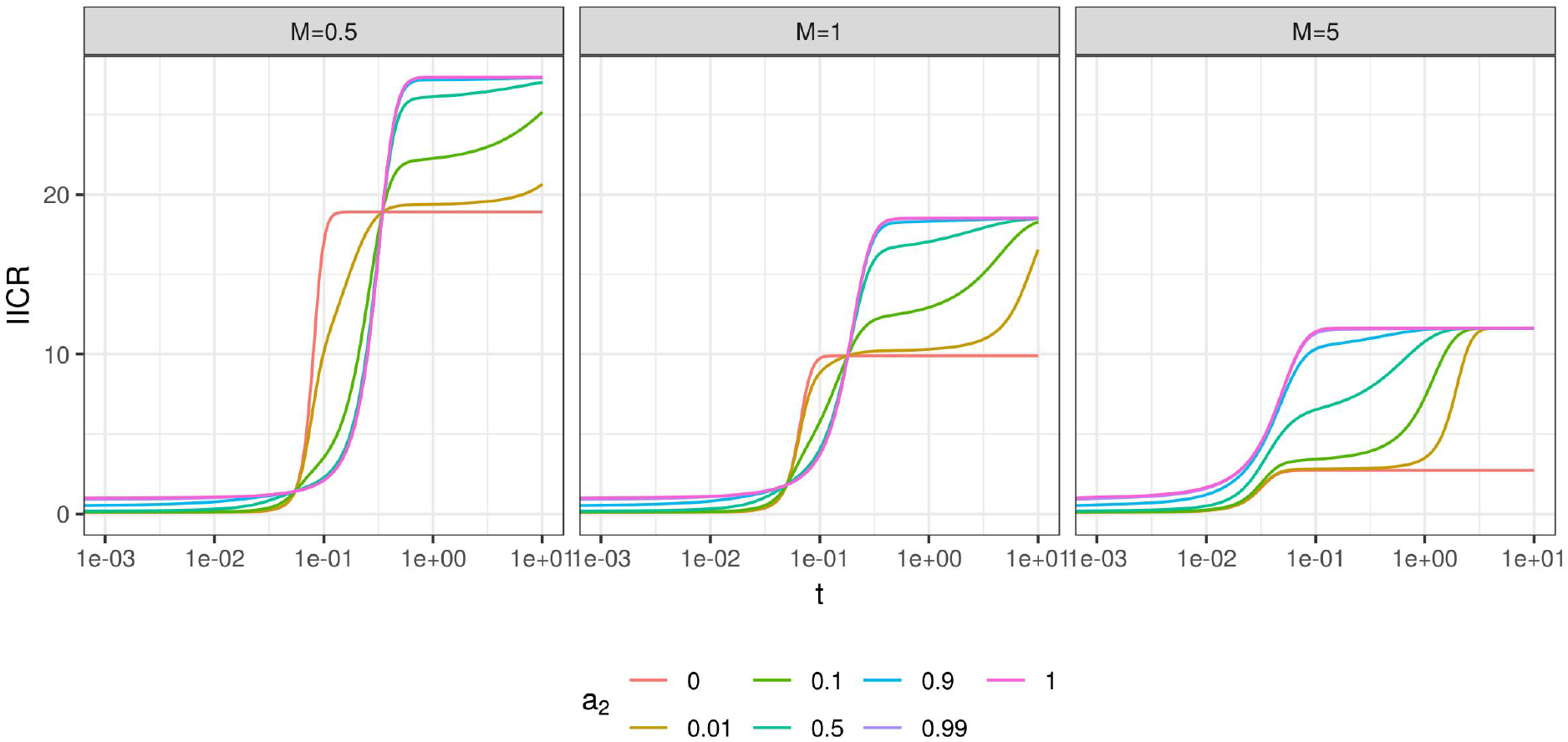
IICR curves for a symmetrical n-island model with *n* = 10 demes and *K* = 2 classes of genomic regions. Regions of class 1 and 2 have a constant deme size 2*Nλ*_1_ and 2*Nλ*_2_ with *λ*_1_ = 0.1 and *λ*_2_ = 1. Scaled migration rate *M* = 4*Nm* is the same for the two classes, each panel corresponding to a different value of this parameter. Frequencies *a*_1_ and *a*_2_ of the 2 classes are given by the legend (having in mind that *a*_1_ + *a*_2_ = 1). For comparison with panmictic models (in particular those in Figure 1), time is scaled by the meta-population size 2*Nn* rather than by the deme size 2*N* as in Equation (6).

For lower migration rates (*M* ≤ 1 in Figure 5), models with rather large values of *a*_1_ are hard to distinguish from the model with *a*_1_=0 (no selection). For instance, the IICR with *a*_2_ = *a*_1_ = 0.5 is not very different from that with *a*_2_ = 1, in contrast to Figure 1 where panmixia was assumed. This suggests that population structure may tend to mask the effect of positive or negative selection even when a quite important part of the genome is under selection. On the other hand, the IICR with *a*_2_ = 0.01 is more similar to that with *a*_2_ = 0 than under panmixia. This suggests that, in the presence of population structure, models with pervasive selection (99% of the genome with *λ* = 0.1) may be interpreted as neutral models with small effective size (100% of the genome with *λ* = 0.1).

Another interesting observation from Figure 5 is the existence of a time window where the IICR is lower when *a*_2_, corresponding to the largest *N_e_*, is largest, i.e. the IICR is lower for models with a smaller part of their genome under selection reducing *N_e_*. This time window occurs in the recent past and is wider for lower migration rates. This counterintuitive result illustrates the limits of interpreting the IICR as a trajectory of effective size, as already outlined for several other demographic scenarios (Chikhi et al., 2018, Mazet et al., 2016). Outside this period, the IICR curves seem to always reach higher values when *a*_2_ is larger. This is in particular the case for *t* close to 0, which is expected analytically (Equation (3)).

### Results: non stationary population structure

To check whether these conclusions may still hold for more realistic evolutionary scenarios, we next assume that each genomic class evolves under the non stationary n-island model estimated by Arredondo et al. (2021) to fit the observed PSMC of a modern human from Karitiana (Li and Durbin, 2011). This model includes 11 islands with symmetric migration and (diploid) deme size 1,380 and it assumes that these islands go through 4 changes of connectivity in the past: *M* ≈ 0.9 (*m* ≈ 1.6e-4) from present to 24,437 generations before present (BP), *M* ≈ 17.7 (*m* ≈ 3.2e-3) from 24,437 to 82,969 generations BP, *M* ≈ 2.5 (*m* ≈ 4.5e-4) from 82,969 to 107,338 generations BP, *M* ≈ 0.7 (*m* ≈ 1.3e-4) from 107,338 to 179,666 generations BP and *M* ≈ 1.1 (*m* ≈ 2e-4) in more ancient times. We define *K* classes of genomic regions: one neutral region with deme size *N* and *K* − 1 other regions under selection with deme size *λ_i_N*, for *λ_i_* either smaller or larger than 1. Results are shown in Figure 6, where two different options are considered to model the heterogeneity of effective size along the genome: (i) the hypothetical three class model of Figure 2 with one class corresponding to positive or negative selection and one other corresponding to balancing selection (top panels), and (ii) the 25 class model of Figure 3 estimated from Gossmann et al. (2011)’s analysis of human real data (bottom panel).

**Figure 6:**
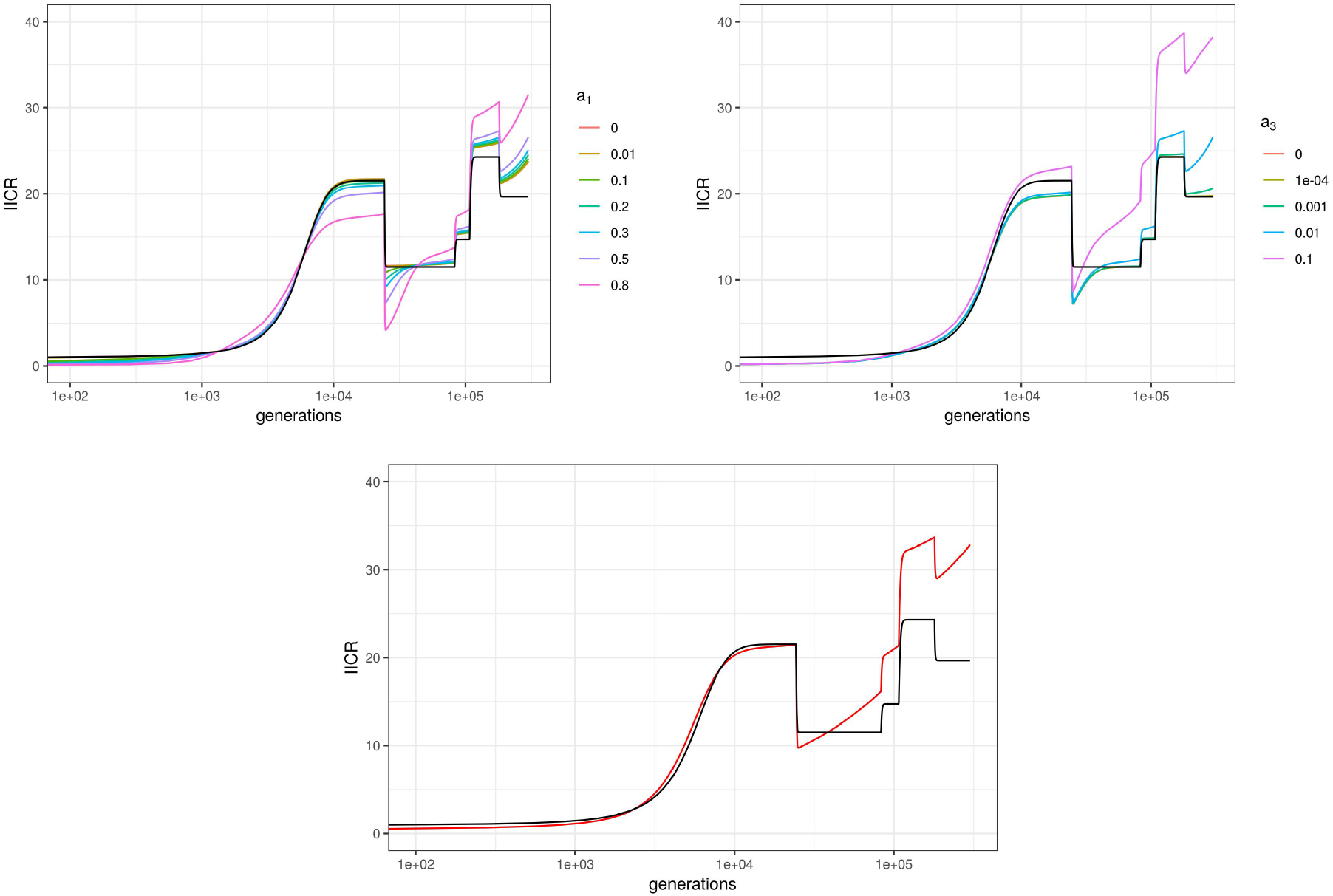
IICRs for demographic models combining population structure and linked selection in humans. The neutral part of the genome evolves under the non stationary n-island model estimated by Arredondo et al. (2021) to fit the observed PSMC of a modern human from Karitiana (Li and Durbin, 2011). This model includes 11 islands with (diploid) deme size *N* = 1380, whose connectivity varied along time according to a 3 step process (see the text for details). To account for selection, this neutral class only represents a fraction of the genome and other classes with lower or higher *N_e_* are also considered. The number of these classes, their proportions and deme sizes (relative to the neutral class) are taken either from Figure 2 (top, where *a*_3_ is fixed to 0.01 in the left panel, and *a*_1_ fixed to 0.5 in the right one) or from Figure 3 (bottom, red line). The black curve on all panels depicts the IICR for this demographic scenario but without selection. Time is shown in generations and in log10 scale.

We find that large values of *a*_1_ could have a significant impact on the IICR in the period ranging from 10,000 to 30,000 generations ago (corresponding to 200-300,000 to 600-900,000 years ago). For instance with *a*_1_ = 0.8, the IICR is around 17 in the most recent hump and around 5 in the most recent “valley”, versus 22 and 12 without selection (top left panel). However, this effect is very moderate when considering the *λ_i_* distribution estimated by Gossmann et al. (2011) (bottom panel). Much more dramatic is the effect observed in the ancient past above 100,000 generations (≈ 2-3 million years) before present, where the IICR with selection is significantly larger than the neutral IICR. This difference is driven by the part of the genome with large effective size (i.e. under balancing selection) and is found (with varying magnitude) in all scenarios.

While the neutral model considered here was estimated without accounting for selection and may thus be itself a biased representation of the true neutral history, the results shown in Figure 6 provide a first approximation of the impact of linked selection on demographic inference in a realistic scenario.

### Methods: modelling transient selection

We finally apply this general framework to model the transient effect of recent selective sweeps, rather than the effect of recurrent positive, negative or balancing selection considered until now. For this analysis we consider a panmictic population. A similar question was tackled by Schrider et al. (2016), who showed in their Figure 5 the estimations obtained when applying the PSMC to a 15Mb genomic region that experienced one or several recent selective sweeps. We focus here on a scenario similar to theirs, with one single selective sweep and approximate the resulting IICR using a model with different classes of *λ_i_* that are time-dependent. In contrast to the model considered in Figure 4, these temporal variations differ between classes, because they depend on the distance to the selected site. Although this model is built based on the expected variations of effective size (or coalescence rate) in a 15Mb region, we note that it also applies to a whole genome having experienced on average one recent selective sweep per 15 Mb region. In other words, our aim here is not to switch from the analysis of global to local IICRs, but rather to explore the local and implicitly global effects in a relatively realistic example.

To approximate the IICR resulting from a recent selective sweep, we assume that the effect of this sweep can be modelled by a reduction of effective population size that is limited both in time (from the emergence of the derived favorable allele to its eventual fixation in the population) and in “genomic space” (i.e. in a genomic neighborhood of this selected variant). More precisely, we consider that the region affected by the sweep on one side of the selected locus is of size

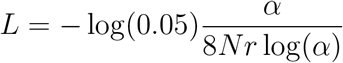

with *N* the diploid population size, *r* the per site recombination rate and *α* = 2*Ns* the scaled selection intensity (*s* being the fitness advantage of homozygotes carrying the selected mutation). This quantity corresponds to the distance in base pairs (bp) from the selected site such that heterozygosity is reduced by only 5% at the end of the sweep (Walsh and Lynch, 2018, chap. 8). To capture the fact that the reduction of effective size caused by the sweep depends on the physical distance to the selected site, we further divide this affected region in 10 classes of size 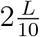 with increasing distance from the sweep, where the factor two results from the sweep extending on both sides of the selected site.

Modelling the selective sweep under the classical “star-like” hypothesis (Nielsen et al., 2005), we approximate (see the Supplementary Material) the average coalescence rate during the sweep as

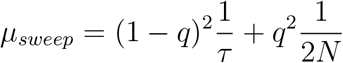

where

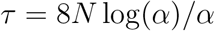

is the duration of the sweep (in generations) and

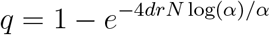

is the per lineage probability of recombination between the selected site and the genomic class. Thus, the relative effective population size in a given genomic class affected by the sweep is equal to 1 before and after the sweep and to

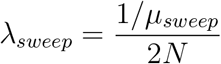

during the *τ* generations of the sweep. A neutral class with *λ* = 1 at all times is also included to account for positions within the 15Mb segment but with physical distance to the selected site greater than *L*.

### Results: transient selection

As shown in Figure 7, top panel, the resulting IICR for *α* = 200 (corresponding to *s* = 0.01 for *N* = 10, 000) is very close to that of a neutral scenario. The IICR for *α* = 1000 (corresponding to *s* = 0.05 for *N* = 10, 000) shows a reduction of about one half at sweep time, similar to the average PSMC plot in Figure 6B of Schrider et al. (2016). The IICR for *α* = 10000 (corresponding to *s* = 0.5 for *N* = 10, 000 or to *s* = 0.05 for *N* = 100, 000) shows a much stronger decline, down to almost zero. However, the IICR decline in our analysis is very localized in time, while the PSMC decline in (Schrider et al., 2016) extends for a longer period. Another important difference is that the PSMC plot in the simulations of Schrider et al. (2016) not only recovers the neutral value after the sweep but increases up to more than twice this value in the recent past. To understand these differences, we simulated coalescence times along a 15Mb region under the same sweep scenario, with *α* = 1000, using the software *msms* (Ewing and Hermisson, 2010) and estimated the resulting empirical IICR as in Chikhi et al. (2018).

**Figure 7:**
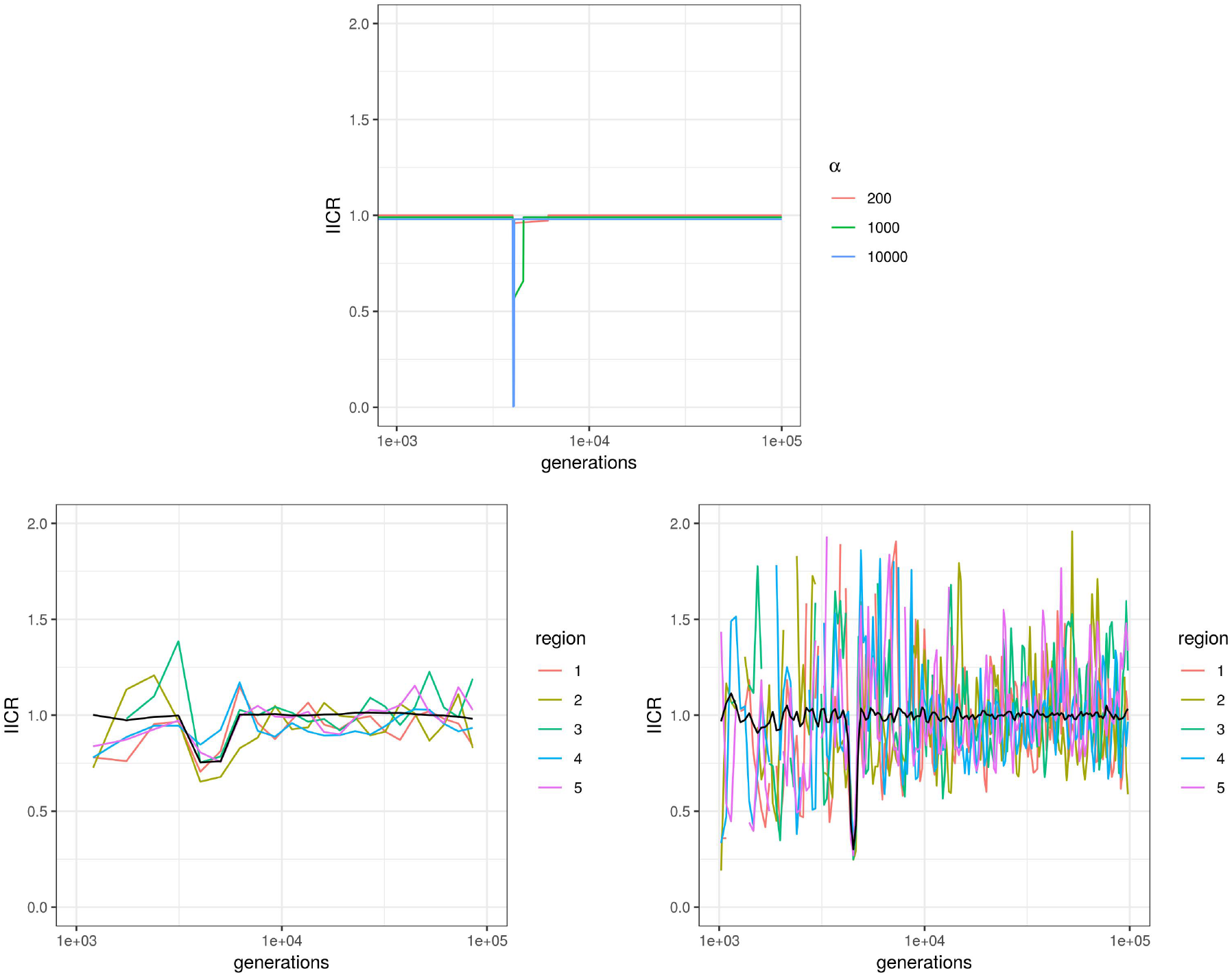
IICRs for a 15Mb region experiencing a single recent selective sweep. Parameter values were chosen to reproduce those in Figure 5 of Schrider et al. (2016): *N* = 10000 (diploid size), *r* = 10^−8^ (per site recombination rate) and *t*_0_ = 4000 generations before present (time where the derived allele got fixed). Times are given in generations and are shown in log10 scale. Top: Expected IICRs when modelling selection using a panmictic model with *K* = 11 classes of regions. Class 11 represents the neutral part of the region (unaffected by the sweep), with relative population size *λ*_11_ = 1. Class *j* (1 ≤ *j* ≤ 10) represents a part of the region affected by the sweep, with a given physical distance from the selected site (which increases with *j*). Relative population size is equal to *λ_j_* = 1 before and after the sweep and is decreased during the sweep to match the larger coalescence rate (see the text for more details). The proportion of each selected class *j* ≥ 10 is *L/*5, where *L* is the size of the region affected by the sweep on either side of the selected site. Scaled selection intensity *α* = 2*Ns* was equal to 200, 1000 or 10000 (see the legend). Bottom: Empirical IICRs based on coalescence times simulated with the software *msms*, for *α* = 1000. Two hundreds independent 15Mb regions were simulated. Colored lines show the IICRs for 5 of these regions (taken at random) and thus represent typical local IICRs. Black lines show the IICRs obtained when merging coalescence times from all regions, they thus correspond to genome-wide IICRs obtained for a 3Gb genome (200 × 15Mb) with one selective sweep every 15Mb. The number of time windows considered (i.e. of distinct estimated IICR values) was equal to 25 (left) or 200 (right) and the length of these windows was increasing exponentially backward in time, as in the PSMC approach.

Similar to PSMC estimations, these empirical IICR estimations depend on the number of time windows considered, the assumption being that *N_e_* is constant within each time window but may vary between time windows. In the bottom left panel of Figure 7, we consider 25 time windows, which corresponds to the order of magnitude used in most PSMC studies. The resulting IICR, averaged over 200 replicates, is transiently reduced around the sweep time and shows no increase above 1 in the recent past, similar to our theoretical prediction (top panel). However, the reduction of *N_e_* is both longer and of lower magnitude than in our prediction, as in the PSMC plots of Schrider et al. (2016). In the bottom right panel, we consider 200 time windows and obtain an average IICR in which the magnitude and duration of the decrease is much more consistent with our theoretical prediction. IICRs from single replicates also correctly capture this reduction around the sweep time but are very noisy outside this period as a side effect of the finer time discretization. Altogether, these results show that modelling selective sweeps by local transient changes of population size leads to a reasonable approximation of the IICR (or equivalently of the genome-wide distribution of *T*_2_) but that discretizing time using a limited number of time windows may lead to soften the true sweep signature by an averaging effect. They also outline that some aspects of a PSMC estimation, as the recent expansion following the sweep in the study of Schrider et al. (2016), cannot be predicted by the IICR, whatever method is used to compute the IICR. The next section explores in more details the link between IICR predictions and PSMC estimations.

## IICR predictions and PSMC estimations

The models and results presented so far allow to predict the effect of linked selection on the IICR, or equivalently on the genome-wide distribution of pairwise coalesnce times.

However, coalescence times are not directly observed from real data so the IICR is in practice estimated from methods like PSMC or MSMC. When population size history is homogeneous along the genome (i.e. *K* = 1 class), PSMC generally provides a very good estimation of the IICR (Mazet et al., 2016) (taking apart considerations relative the amount or the quality of the data). But when population size history is heterogeneous along the genome, as considered here to approximate the effects of selection, the answer may depend on the scale (10kb? 100kb? 1Mb?) at which this heterogeneity is detectable. In other words, for a fixed proportion of genomic positions with reduced effective size due to linked selection, PSMC results may depend on the spatial clustering of these positions along the genome, while the IICR does not.

To explore this question, we tested whether genomic data including genome-wide heterogeneity of *N_e_* at different scales could generate PSMC plots consistent with our IICR predictions. To do this we carried out a limited number of additional simulations in which, using the genomic sizes *λ*_1_ = 0.1 and *λ*_2_ = 1, we varied the lengths *L*_1_ and *L*_2_ of contiguous DNA chunks belonging to a given class, while keeping constant the proportions *a*_1_ and *a*_2_ = 1 − *a*_1_ at which these classes are represented. The lengths *L*_2_ for the chunks of class 2 were chosen to be 10^6^, 10^5^ and 10^4^ base pairs, and the lengths for the chunks of class 1 followed from the proportions *a*_1_ and *a*_2_. We tested three values for the frequency *a*_1_ (0.5, 0.9 and 0.99), and for each combination of *a*_1_ and *L*_1_ we simulated two independent genomes of length 10^9^ base pairs, where the two size classes were evenly spaced in the form *L*_1_*, L*_2_*, L*_1_*, L*_2_, …, *L*_1_*, L*_2_. We found that PSMC estimations fit well IICR predictions for large chunks (*L*_2_ = 10^6^ and 10^5^), but may highlight more complex and unpredicted patterns for smaller ones (Figure 8).

**Figure 8:**
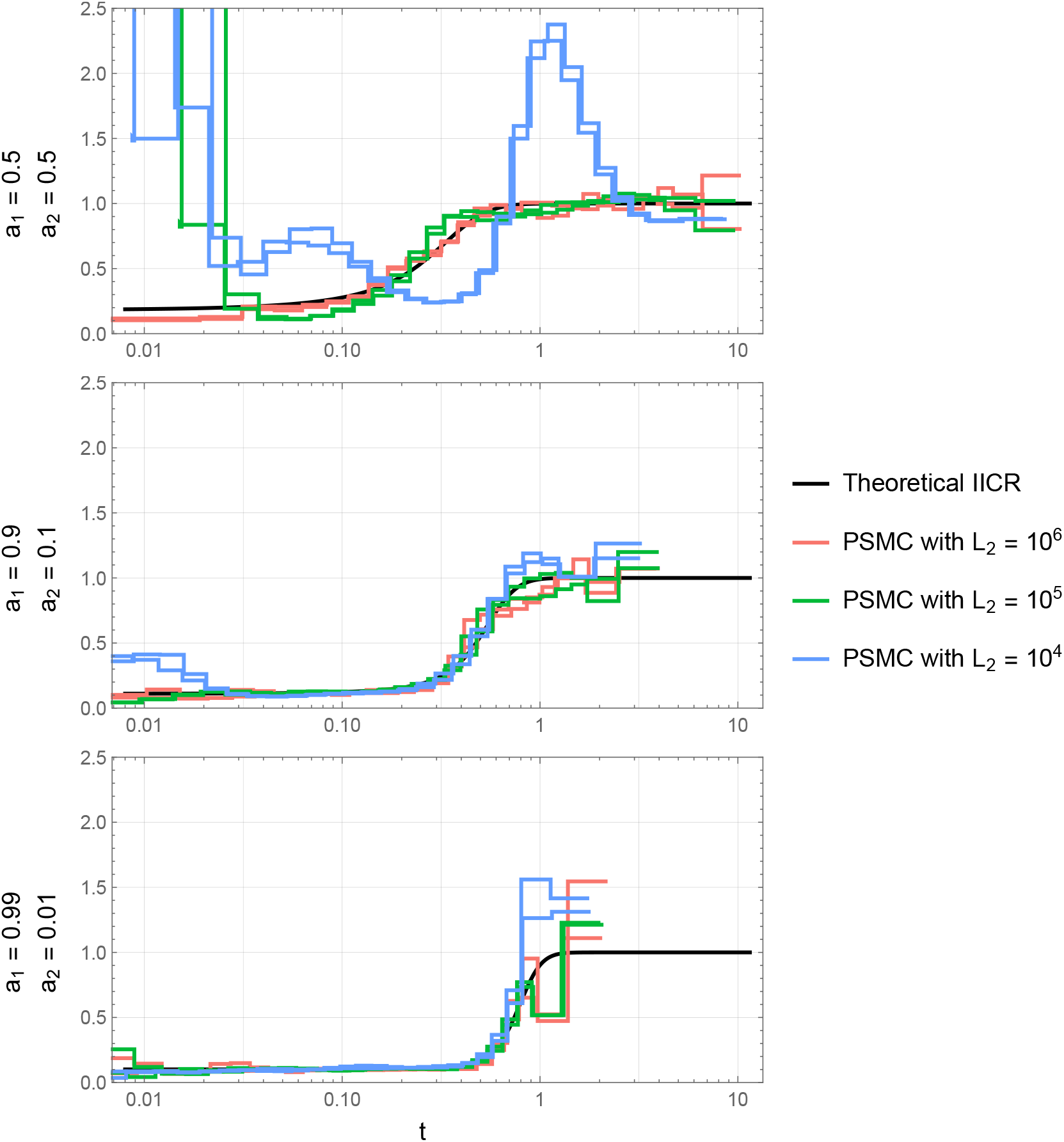
Comparison between theoretical IICR and inferred PSMC. For each frequency distribution (*a*_1_*, a*_2_) of the two size classes *λ*_1_ = 0.1 and *λ*_2_ = 1 we show the corresponding theoretical IICR (black) and two independent PSMC simulations for three values of the chunk length *L*_2_. In each case, 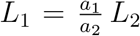. The simulated sequence has a total length of 10^9^ bp and the two class chunks are evenly alternated in the form (*L*_1_*, L*_2_*, L*_1_, …, *L*_2_). Population size was equal to 10000.

## Discussion

### Effects of linked selection on the IICR

A classical assumption in population genetics considers that linked selection can be modelled as a first approximation by a local change in effective population size (Hill and Robertson, 1966). Background selection and selective sweeps, which tend to reduce genetic diversity locally (Charlesworth et al., 1993, Smith and Haigh, 1974), are then seen as resulting in lower *N_e_* values, whereas genomic regions under balancing selection are in contrast interpreted in terms of higher *N_e_* values. In both cases, the impact of selection on genetic diversity or *N_e_* is stronger for regions with lower recombination or higher selective constraints (number of selected sites, selection intensity) (Charlesworth, 2009). At the genome-wide level, linked selection appears thus to generate an apparent heterogeneity of *N_e_* among genomic regions, reflecting the variations of the mode (increasing or decreasing *N_e_*) and the intensity of linked selection (Gossmann et al., 2011, Jiménez-Mena et al., 2016a). Following this simplifying assumption, we described in this study the distribution of the coalescence time between two sequences (*T*_2_) for models including variable classes of *N_e_* along the genome. More precisely, we characterized the IICR (Mazet et al., 2016) of such models, a quantity that is equivalent to the *T*_2_ distribution and corresponds to the graphical output of the popular PSMC approach (Li and Durbin, 2011), which is generally interpreted as the past temporal trajectory of *N_e_* of the population or species under study. This analysis allowed us to predict the expected effects of linked selection on PSMC or related demographic inference approaches (Schiffels and Durbin, 2013).

One of the main conclusions of our work is that, under panmixia and constant population size, the existence of several classes of *N_e_* (induced by linked selection) *always* results in a spurious signal of population size decline: the IICR of such models is a decreasing function (forward in time) whose highest value (reached in the ancient past) corresponds to the largest genomic *N_e_* and lowest value (reached in the most recent past) to the harmonic mean of genomic *N_e_* values weighted by their relative proportion in the genome (Figure 1, Equation 3). Specifically, we found that selection reducing *N_e_* (background selection or sweeps) has a stronger effect on the IICR in the recent past, while selection increasing *N_e_* (balancing selection) mainly influences the IICR in the intermediate and ancient past (Figure 2). There is a striking asymmetry between the two forms of selection: because the IICR plateau is determined by the class with the largest *N_e_* independently of the proportion of this class, even a minute proportion of balancing selection can have a large effect on the IICR, whereas higher proportions of background selection or sweeps are necessary to generate significant and detectable effects on the IICR (Figure 2). Combining the two forms of selection by considering *N_e_* distributions inferred from real data (Elyashiv et al., 2016, Gossmann et al., 2011) we found that linked selection is expected to cause a long term apparent five-fold decrease of the IICR in organisms such as humans or *Drosophila melanogaster* (Figure 3). However, we stress that these results assumed panmixia and constant population size.

Another important conclusion of our work is indeed that the effects of linked selection on the IICR mentioned above may be largely hidden by those of population structure. Considering a symmetrical *n*-island model, we observed for instance that even when a large proportion of the genome is influenced by selection reducing *N_e_* the effect on the IICR could be difficult to see for models with reduced migration rates between islands (Figure 5). Focusing on humans we also considered a simple but reasonable demographic scenario of variable population structure (Arredondo et al., 2021) together with a realistic genomic *N_e_* distribution for this species (Gossmann et al., 2011). We found that the largest and most visible effect of linked selection on the IICR was an ancient population size decline related to the presence of balancing selection (Figure 6, bottom).

Such ancient declines are indeed observed in PSMC plots inferred in humans and a number of other species, but a further complication is that these patterns may also arise due to the low number of informative coalescence events available to PSMC in this ancient time period. PSMC analyses of genomic data simulated under realistic demographic scenarios, with and without balancing selection, will be necessary to investigate whether these ancient signatures of balancing selection can be disentangled from statistical artifacts. As a simple test we simulated genomic data under the demographic model of Figure 6 with a single genomic *N_e_* (i.e. no selection). We applied PSMC to these data and found no ancient decrease in the estimated trajectory compared to the expected IICR (Figure S5). These admittedly limited results suggest that the PSMC is not necessarily *statistically* biased in the ancient past, and that the signals observed in several species including humans and chimpanzees might be due to balancing selection or other forms of selection maintaining high levels of diversity over very long periods. One possible strategy to limit the influence of regions submitted to such forms of selection would be to first detect them and filter them out from the PSMC analysis. For the demographic scenario of Figure 6, we found that this would reduce the biases observed in the ancient past without affecting significantly other parts of the IICR (Figure S6).

### The intriguing signature of background selection on the IICR

The framework developed in this study makes no particular distinction between positive and background selection, which are both modelled as leading to a reduction of *N_e_*. Thus, one possible interpretation of our results would be that ignoring background selection leads to infer spurious population declines. This conclusion is at odds with several previous studies, which concluded that unaccounted background selection may actually lead to a spurious signature of recent population expansion. For instance, Zeng and Charlesworth (2011) and Walczak et al. (2012) developed theoretical approximations of the genealogical process at a neutral locus linked to a site under negative selection and showed that this process shared many properties with that of an expanding population. The former study accounted for intra-locus recombination, whereas the latter ignored it. Several recent studies have applied demographic inference methods to genomic data simulated with and without background selection (Ewing and Jensen, 2016, Johri et al., 2021, Lapierre et al., 2016, Pouyet et al., 2018) and observed a signal of recent population expansion in the scenarios including selection. Finally, Johri et al. (2020) analyzed real data from an African population of *Drosophila melanogaster* with a new ABC demographic inference approach accounting for background selection. They estimated that the size of this population has been relatively constant for a few millions generations, while several previous studies on this or other related populations, which ignored background selection, estimated a strong recent population size increase, e.g. (Arguello et al., 2019, Kapopoulou et al., 2018).

Two main reasons may resolve this apparent paradox between these previous results and ours. First, we assume that linked selection can be modelled by a local change of *N_e_* without any temporal dynamics (except in Figure 7 and related text, whose focus is specifically on recent selective sweeps). In particular, our results do not hold for demographic inference approaches based on the Site Frequency Spectrum (SFS), because weak background selection is expected to produce an excess of low frequency alleles, in particular singletons, which cannot be mimicked by just assuming a smaller *N_e_*. Such an excess of rare alleles is also a classical signature of expanding populations, which may explain the conclusions of several of the studies mentioned above (Ewing and Jensen, 2016, Johri et al., 2020, Lapierre et al., 2016, Pouyet et al., 2018).

Second, even when focusing on pairwise statistics such as heterozygosity or *T*_2_, the signature of population decline predicted by the IICR can only be observed if the data considered exhibit some heterogeneity in *N_e_*. As it can easily be seen from Figure 1, panmictic models with either no (*a*_2_ = 1) or only (*a*_2_ = 0) selection do not show declining but constant IICRs. Consequently, a decline signature is not necessarily expected when analyzing a single locus under selection as in Zeng and Charlesworth (2011) or Walczak et al. (2012). It is also not necessarily expected when analyzing genome-wide data with homogeneous selective constraints along the genome. For instance, Johri et al. (2021) simulated genome-wide sequences including background selection by considering a regular alternance of functional (selected) and intergenic (neutral) regions of fixed and relatively small sizes: depending on the scenario, the size of a single ‘unit’ including one functional and one intergenic region ranged from ≈ 13 to 55 kb. The PSMC analyses of these sequences suggested a population under constant size or slight recent expansion. We believe that some of the results obtained by these (and possibly other) authors could be due to the fact that the data simulated with this approach do not exhibit enough heterogeneity in population sizes among (short) sliding windows over the genome. Such a regularity is at odds with observations made in different organisms (Elyashiv et al., 2016, Gossmann et al., 2011).

### IICR predictions and PSMC estimations

Understanding the difference between our results and those of Johri et al. (2021) also leads to the fundamental question of the link between a PSMC curve and the IICR. The results obtained in Figure 8 suggest that the IICRs computed in this study are good predictors of PSMC outputs when variations of *N_e_* occur at a relatively large scale (100 kb or more), but not always when these variations occur at a smaller scale. This may explain the discrepancy between our predictions and the PSMC results in the scenario simulated by Johri et al. (2021), where the heterogeneity of *N_e_* was detectable only at very small scale (≤ 55kb).

The recent selective sweep scenario considered in Figure 7 provides another example of potential differences between PSMC estimations and IICR predictions in the case of genomic heterogeneity. Simulating *genome sequences* in a single 15Mb region experiencing one recent selective sweep, Schrider et al. (2016) found that PSMC applied to these sequences would infer a bottleneck around the time of the sweep completion, generally followed by a more recent expansion exceeding the ‘neutral’ effective size. Simulating *coalescence times* under the same selective sweep scenario and estimating the IICR from these simulated values, we observed a similar bottleneck but no recent expansion. This difference likely results from the fact that short coalescence times are mostly clustered around the selected site in the real data, while for IICR estimation only their proportion over the 15Mb region matters. Approximating the IICR under a selective sweep through a model with several classes of time-dependent *N_e_*, we managed to reproduce the main characteristics of the IICR of this scenario, but this is not exactly similar to the PSMC that would be estimated in this scenario.

Overall, these results suggest that assessing potential PSMC biases in a given species may require specific simulations based on precise genomic annotations (positions and lengths of genes, local recombination rates …). As an alternative to such specific studies, we provide here a quick and flexible approach to predict the distribution of coalescence times in the presence of linked selection, which is to some extent also representative of expected PSMC outputs.

### Perspectives for demographic inference

The above discussion illustrates that the effects of linked selection on demographic inference are complex, as they not only depend on the type and intensity of linked selection but also on the inference approach applied (SFS or *T*_2_ based for instance) or the scale at which selection constraints vary along the genome. If the future confirms that linked selection is pervasive in the genome as claimed for several model species (Elyashiv et al., 2016, Pouyet et al., 2018) new demographic inference approaches accounting for linked selection and population structure will be needed. One way of achieving this objective is to jointly estimate demographic and selection parameters, as proposed in two recent studies relying on simulation based approaches, deep learning (Sheehan and Song, 2016) and Approximate Bayesian Computation (ABC) (Johri et al., 2020). These studies focused on relatively simple models, considering panmictic populations with a single population size change and only some types of selection (background selection in one study, sweeps and balancing selection in the other). To integrate more complex demographic scenarios, several recent studies considered demographic models including two classes of *N_e_* along the genome, one for neutral loci and one for loci under linked selection. The proportion of the two classes and the ratio of *N_e_* between them were estimated together with other parameters of the demographic model, using either ABC (Rougemont and Bernatchez, 2018, Roux et al., 2016) or a modification (Rougemont et al., 2020, Rougeux et al., 2017) of the diffusion approach implemented in the software *∂*a*∂*i (Gutenkunst et al., 2009). Our study suggests that a similar inference approach, accounting for linked selection through variable classes of *N_e_* along the genome, could be developed based on the IICR. An IICR-based inference framework was recently proposed for the estimation of non stationary *n*-island models and provided very encouraging results (Arredondo et al., 2021). Given the strong impact of linked selection on the IICR under panmixia, we believe that a similar approach could allow to jointly infer parameters related to demographic history and to the *N_e_* distribution. However, the results obtained under models of population structure suggest that it may be necessary to use the IICR in addition to other summaries of genomic diversity to overcome identifiability issues. Also, we should stress that separating the effects of population size change, selection and population structure is likely to be one of the major challenges of population genetics in the future.

### Pros and cons of an IICR approach

Whether the objective is to predict potential effects of linked selection or to estimate linked selection parameters from real data, two nice features of an IICR-based approach such as the one considered here are flexibility and speed of computation. This approach allows to simultaneously include different forms of selection and to combine linked selection with arbitrary complex demographic models. The examples considered here included for instance panmictic models with temporal variations of the population size (Figure 4) and n-island models with temporal variations of the migration rate (Figure 6). We also considered different distributions of *λ_i_*, some of them including a large number of classes. More general models could be considered, for instance including other forms of structure or combining population structure and temporal population size variations. In the case of structured models, variable migration rates along the genome may be considered: we could either decrease *M* in the linked selection class(es) to account for possible effects of selection on migration success or introduce new classes with lower *M* values in order to model possible barriers to gene flow (Roux et al., 2016). As outlined in Figure 7, transient selection can be modelled by including population size changes in a subset of classes, and this approach could also be extended to model more complex fluctuating selection effects. Whatever the complexity of the demographic model and the *N_e_* distribution considered, the associated IICR can be computed exactly in a very small time using the rate matrix approach described in Rodríguez et al. (2018) or Arredondo et al. (2021), which allows to efficiently explore a very large number of scenarios or parameter values.

We should also stress that apparent variations of *N_e_* along the genome may result from other biological processes than linked selection. The models presented here, and the general conclusion that heterogeneity in *N_e_* is expected to generate population size decline patterns, also apply to these other biological processes. For instance, genomewide variations of the mutation rate may have similar effects on the data than genomewide variations of *N_e_*, because high mutation rates and large population sizes both lead to increase the number of polymorphic sites in a region. Consistent with our results, Sellinger et al. (2021) showed that applying SMC methods to genomic sequences that were simulated with local variations of the mutation rate leads to infer spurious population size declines. Actually, a direct consequence of *N_e_* heterogeneity is to increase the variance of coalescence times along the genome (see the Supplementary Materials for a proof of this statement under panmixia). Inference methods like the PSMC, which do not account for *genomic* variations of *N_e_*, try to explain this additional variance using *temporal* variations of *N_e_*, more precisely population size declines.

The main limitation of the IICR approach described in this study is that it focuses on pairs of sequences. It provides information that is complementary to that provided by the SFS, as we have noted elsewhere (Arredondo et al., 2021, Chikhi et al., 2018) For instance, some effects of weak background selection or selective sweeps may be visible on the SFS but not on the IICR. Currently we have mainly focused on the IICR as defined for a pair of sequences, but extensions to multiple sequences might provide additional information on the distribution of higher order coalescence times (*T*_3_, *T*_4_, …)., hence allowing a finer characterization of selective and neutral processes.

### Closing comments

We have used the IICR as a way to explore important ideas that are central to population genetics such as the notion of effective size (see also Chikhi et al. (2018), Mazet et al. (2016) for discussions on these questions), drift and selection. We wished to re-open discussions regarding the influence of selective and neutral processes on genetic diversity, some of them general and theoretical, others more specific and practical: Can selection be modelled as a genomic variation in *N_e_*? What are the limits of such an approximation? Can linked selection, and more generally *N_e_* variation along the genome, be detected in real genomes by applying the PSMC method of (Li and Durbin, 2011) or related approaches? These are exciting questions to ask and the recent years have shown that they are at the heart of modern population genetics.

## Data availability statement

Code used to generate the exact and simulated IICRs shown in this study can be found at https://github.com/sboitard/IICR_selection.

## Acknowledgements

Armando Arredondo was funded by the Université Fédérale Toulouse Midi Pyrénées (UFTMiP) and the Région Occitanie (formerly Midi-Pyrénées) with PhD grant No. 31I2017M248. Lounès Chikhi was funded by Fundação para a Ciência e Tecnologia (ref. PTDC-BIA-EVL/30815/2017). Olivier Mazet and Lounès Chikhi were funded by the 2015–2016 BiodivERsA COFUND call for research proposals, with the national funders ANR (ANR-16-EBI3-0014) and the Fundação para a Ciência e Tecnologia ref. Bio-diversa/0003/2015 and PT-DLR (01LC1617A). This work was also supported by the LABEX entitled TULIP (ANR-10-LABX-41 and ANR-11-IDEX-0002-02) as well as the LIA BEEG-B (Laboratoire International Associé-Bioinformatics, Ecology, Evolution, Genomics and Behaviour). We acknowledge an Investissement d’Avenir grant of the Agence Nationale de la Recherche (CEBA: ANR-10-LABX-25-01).

## Supplementary Material

### Monotony of the IICR in a panmictic model with several classes of constant *N_e_*

We consider here the first model introduced in this study, where a proportion *a_i_* of the genome evolves under a Wright-Fisher model with constant population size *λ_i_N* (i=1,…, K). The IICR under this model is given by equation (2). To characterize the dynamics of the IICR over time, we study the derivative of the IICR as a function of time (backward from present):

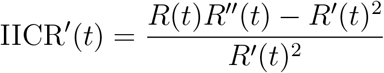

which has the sign of

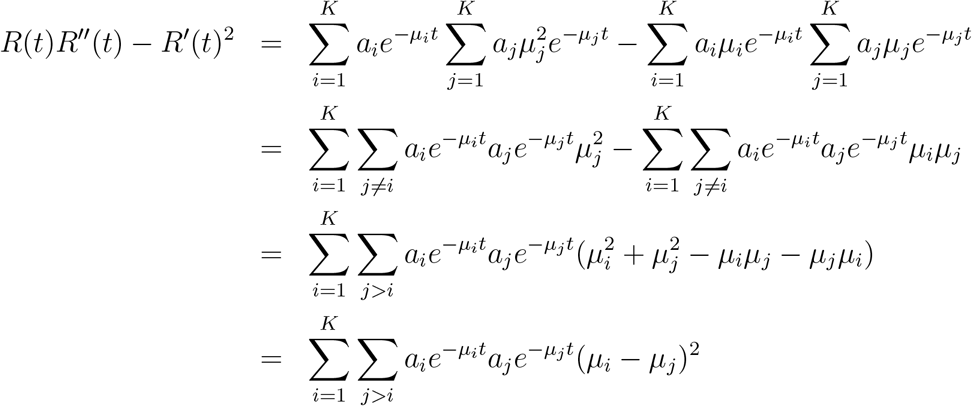

This quantity is always positive so we can conclude that the IICR is *always increasing* from *t* = 0 to *t* = +∞ (i.e. backward in time).

### Variance of *T*_2_ in a panmictic model with several classes of constant *N_e_*

We consider here the same model as in previous section. For a given position in the genome, let us denote *T*_2_ the pairwise coalescence time (in 2*N* units) and *X* the genomic class. *X* is a stochastic variable that is equal to *i* with probability *a_i_*, and the distribution of *T*_2_ conditional on *X* = *i* is an exponential distribution with parameter 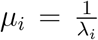. In particular, we have 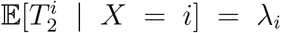 and 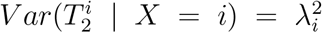. From these assumptions, we can deduce that

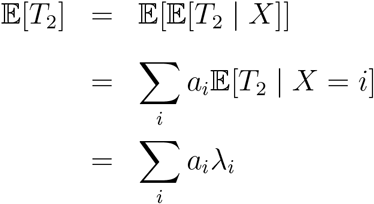

and

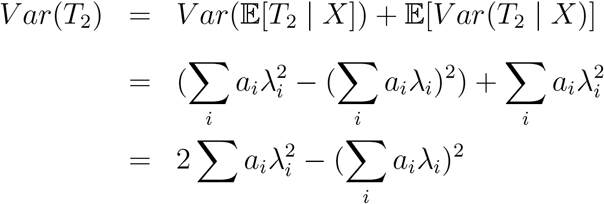

where the derivation from the first to the second line follows from the fact that (i) 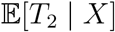 is a stochastic variable equal to *λ_i_* with probability *a_i_* and (ii) *V ar*(*T*_2_ | *X*) is a stochastic variable equal to 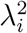 with probability *a_i_*.

In comparison, the variance of *T*_2_ in a model with a single class of *N_e_* and the same expected value of *T*_2_ is

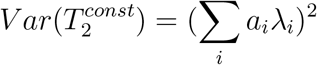

Thus, we have

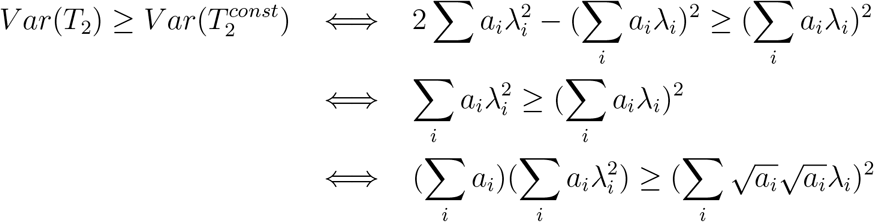

which is always true from the Cauchy Schwartz inequality.

Let us denote 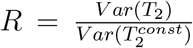 the ratio of the two variances, which is thus always larger than 1. We observed that this ratio generally increased with the proportion of the genome associated to the smallest *λ_i_*. For instance, in the two class model of Figure 1 with *λ*_1_ = 0.1 and *λ*_2_ = 1, *R* was equal to 1.08 for *a*_1_ = 0.1, 1.53 for *a*_1_ = 0.5 and 2.24 for *a*_1_ = 0.1. In the three class model of Figure 2 with *λ*_1_ = 0.1, *λ*_2_ = 1, *λ*_3_ = 3 and *a*_3_ = 0.01 (left panel), *R* was equal to 1.13 for *a*_1_ = 0.1, 1.61 for *a*_1_ = 0.5 and 2.75 for *a*_1_ = 0.1.

### Estimation of the distribution of *N_e_* in drosophila and humans

Two different distributions of *λ_i_* over the genome were obtained for *Drosophila melanogaster*. The first one was taken from the study of Elyashiv et al. (2016), who developed a method for inferring the distribution of fitness effects in different classes of functional annotations (UTRs, codons …) for both beneficial and deleterious mutations. This method requires polymorphism data from the focal species, divergence data with closely related species and precise recombination and annotation maps allowing to assess the selection constraints acting on each position in the genome. A by-product of their analysis is that an estimation of *N_e_* can be obtained for sliding windows along the genome. Interestingly, these *N_e_* values resulting from the strength of linked selection in each genomic region are defined as the inverse of the coalescence rate between two sequences and all computations rely on heterozygosity values observed between pairs of individuals. This suggests that the *N_e_* estimates should be directly comparable with our *λ_i_* values, which also correspond to the inverse of pairwise coalescence rates. The values of *N_e_* estimated by Elyashiv et al. (2016) for 1Mb sliding windows in *Drosophila melanogaster*, based on 162 inbred lines derived from the Raleigh, North Carolina population, were downloaded at *https://github.com/sellalab/LinkedSelectionMaps*. Their distribution (top left panel) was converted into a discrete distribution of *λ_i_* values with *K* = 25 classes using the *hist*() function of R. The IICR resulting from this distribution is shown in the top middle and right panels.

The second distribution used for this species was that estimated by Gossmann et al. (2011) for a Zimbabwe population. While these authors also used polymorphism and divergence data, they focused on exons and did not aim at modelling the distribution of fitness effects. They assumed a log-normal distribution of *N_e_* with mean value of 1 and estimated the scale parameter of this distribution from the observed data at several independent genes in the genome. Using the parameter obtained by this approach for *Drosophila melanogaster* and no recombination within genes (Table 1 of their study), we randomly sampled 100,000 values of *N_e_* (or *λ*) under the log-normal distribution (middle left panel). A discrete distribution of the *λ_i_*’s and the associated IICR were then computed as explained above, filtering out large *λ* values (we arbitrarily excluded values above five). Indeed, it is not clear whether such large values would be realistic or statistical artifacts resulting from the use of a continuous distribution estimated mainly from smaller *λ* values. Also, they represent less than O.6% of the distribution. As a comparison with another species, we also applied this second approach with the scale parameter inferred by Gossmann et al. (2011) for humans based on data from the Yoruba population (bottom panels).

### Derivation of the pdf of *T*_2_ in a *n*-island model

We derive here the pdf density of *T*_2_, the coalescence time of two lineages sampled in the same deme (resp. different deme), in an *n*-island model. We follow the identity by descent approach used in Durrett’s process (Durrett, 2008, p. 150). The size of each deme is *λN*, the probability of each lineage to migrate from a deme to another each generation is *m*, and the per locus mutation rate is *u*. Define the rescaled mutation and migration rates by *θ* = 4*Nu* and *M* = 4*Nm*. Note that two lineages coalesce at rate 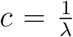 when they are in the same deme, migrate at rate 2*m.*2*N* = *M* and experience mutations at rate 2*u.*2*N* = *θ*.

Let *p_s_*(*θ*) and *p_d_*(*θ*) be the probabilities that two lineages are identical by descent when they are chosen in the same or different demes. Following back two lineages from the same deme, three different events can occur: a coalescence with probability 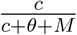, a migration with probability 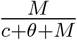 and a mutation with probability 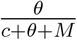. If lineages are in different demes, the only possible events are mutation, with probability 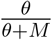 and migration. In this second case lineages arrive in the same deme with probability 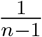 and stay in different ones with probability 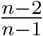. Hence we have the two coupled equations:

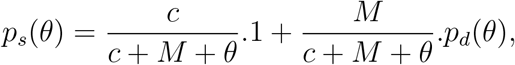

and

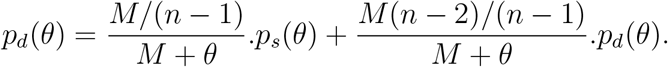

The second equation gives

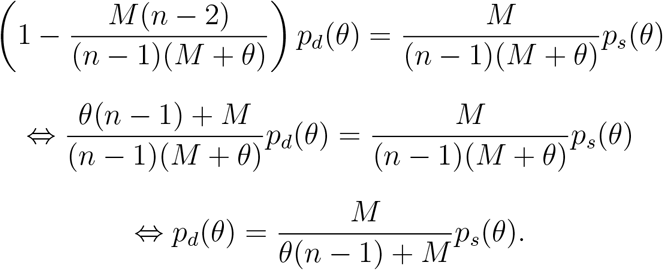

We then inject in the first equation:

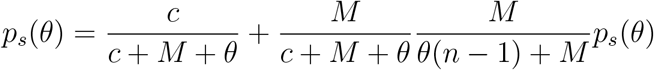

hence

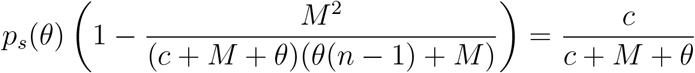

and since

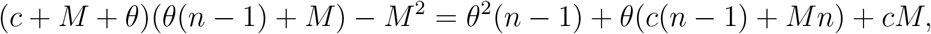

we get

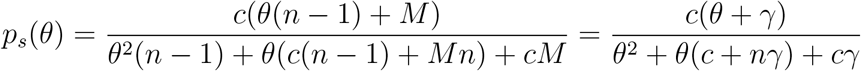

and

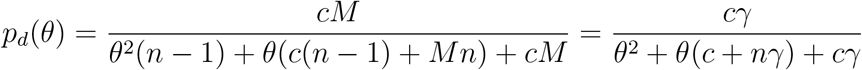

with

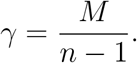

Let’s now note that the probability *p_s_*(*θ*) that two lineages has reached their common ancestor without undergoing any mutation is also the expected value 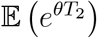. In other words, *p_s_* is the Laplace transform of *T*_2_. It can be inverted by looking for the roots of *θ*^2^ + *θ*(*c* + *nγ*) + *cγ*. Let Δ = (*c* + *nγ*)^2^ − 4*cγ*, then

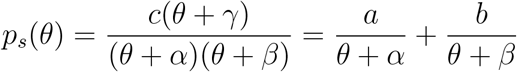

with

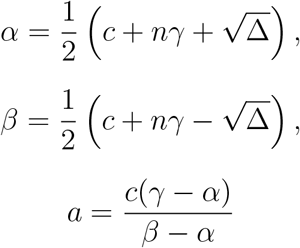

and

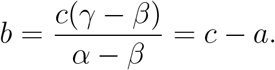

Hence the probability density function of *T*_2_ is:

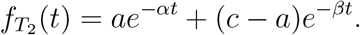

Note that −*α* and −*β* are the non zero eigenvalues of the Q-matrix, −*β* being the closest to 0, and we have the relationships *α* + *β* = *c* + *nγ* and *αβ* = *cγ*. Note also that we could similarly obtain the pdf distribution of the coalescence time of two lineages sampled in different demes, as *p_d_* is its Laplace transform as well.

### Approximation of the coalescence rate in a selective sweep scenario

Assuming a selectice sweep scenario with scaled selection intensity *α*, we consider here the genealogy at a neutral locus located *d* bp away from the selected site. This process can be modelled using a structured coalescent where lineages are either in the ‘derived’ or ‘ancestral’ background, depending on which allele at the selected locus they are associated with (to avoid any confusion, we remind here that this structure is a modelling facility and has nothing to do with the island structure considered in some sections of the main text). In this framework, ancestral recombination events creating or breaking the association with the derived allele can be seen as migration events from one background to the other (Kaplan et al., 1988). In the case of a complete selective sweep, lineages sampled at present all belong to the derived background, because the derived allele is then fixed in the population. Following previous studies on this topic, e.g. (Nielsen et al., 2005), we further assume a “star-like” model where these lineages can either (i) escape this derived background through recombination and stay in the ancestral background until the end of the sweep phase (i.e. at the time when the derived allele appeared, as we go backward in time) or (ii) coalesce all together at the end of the sweep phase. Actually, we slightly relax this second hypothesis and simply assume that their average coalescence time corresponds to the end of the sweep phase. The probability for each lineage to escape the sweep is approximately

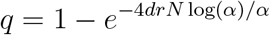

where *r* is the recombination rate per generation and per bp. Because lineages can only coalesce if they are in the same background (derived with probability (1 − *q*)^2^ or ancestralwith probability *q*^2^), we assume that the average coalescence rate during the sweep is

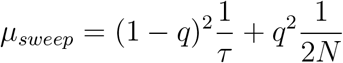

where

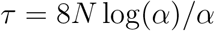

is the duration of the sweep (in generations). In this formula, 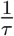 approximates the average coalescence rate for two lineages not escaping the sweep, which follows from our assumption that the average coalescence time is *τ*, and 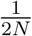 is the standard neutral coalescence rate which applies to two lineages having escaped the sweep.

### Supplementary figures

**Figure S1:**
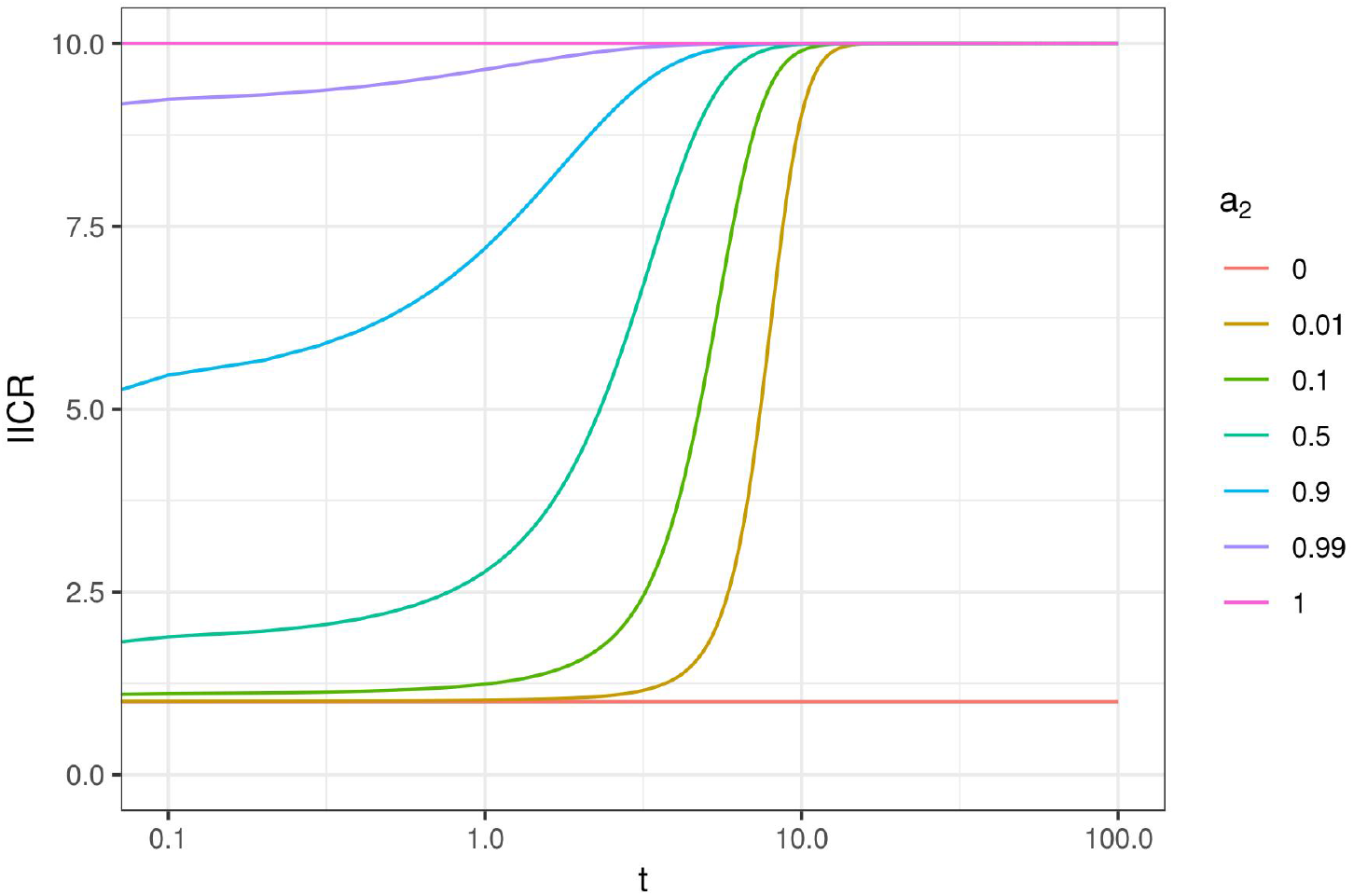
IICR curves for a panmictic model with *K* = 2 classes of genomic regions with constant size. Same as Figure 1 with *λ*_1_ = 1, *λ*_2_ = 10 and time from 0 to 100 (in log10 scale)

**Figure S2:**
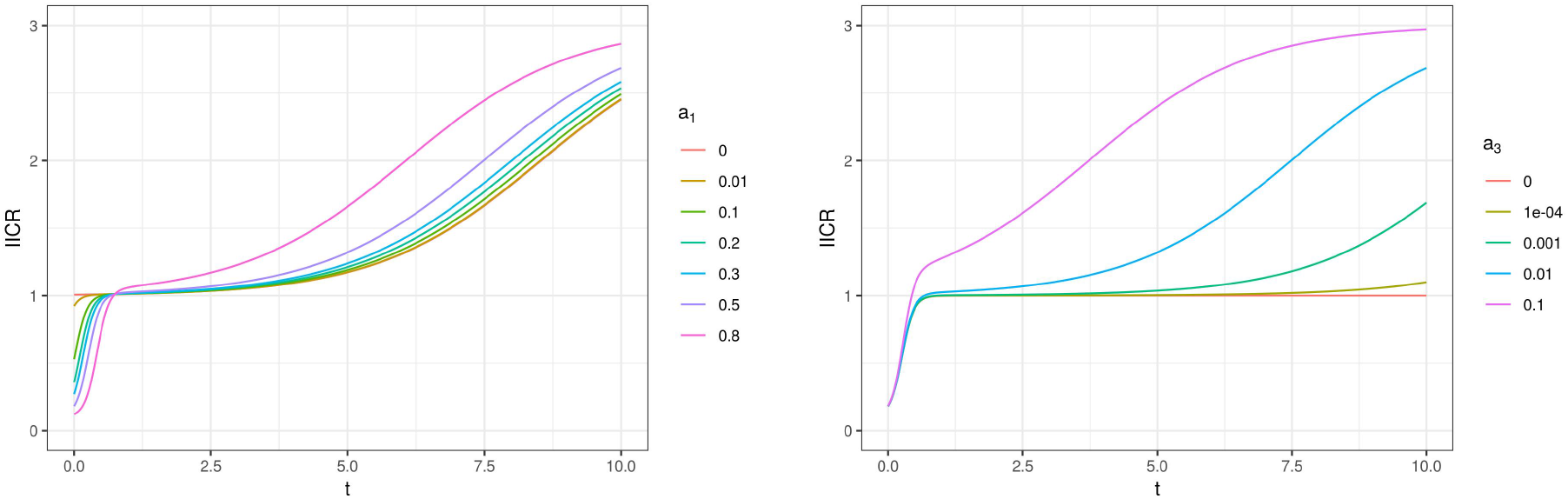
IICR for a panmictic model with *K* = 3 *λ_i_* values such that *λ*_1_ < 1, *λ*_2_ = 1 and *λ*_3_ > 1. Same as Figure 2 except that time is plotted in natural scale.

**Figure S3:**
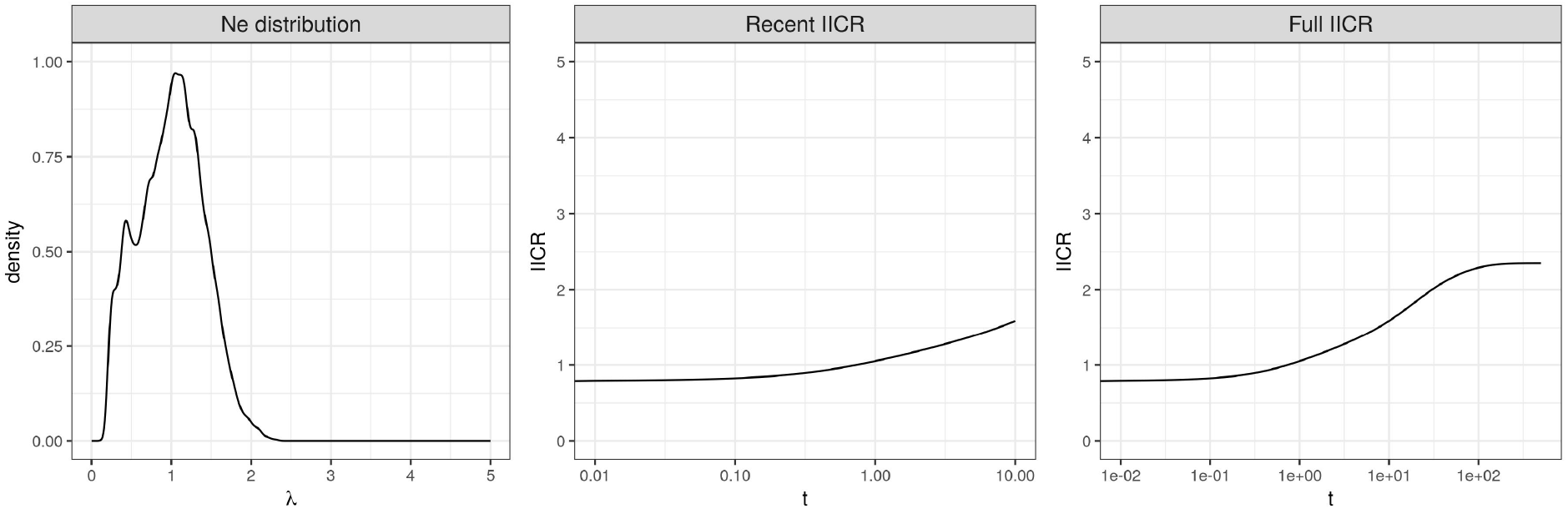
IICR obtained when removing low *N_e_* values from the distribution estimated by Elyashiv et al. (2016). This truncated distribution (rescaled to have a mean of 1 as the others) is shown on the left panel. The associated IICR is shown until *t* = 10 (middle panel) or *t* = 500 (right panel), in log10 scale.

**Figure S4:**
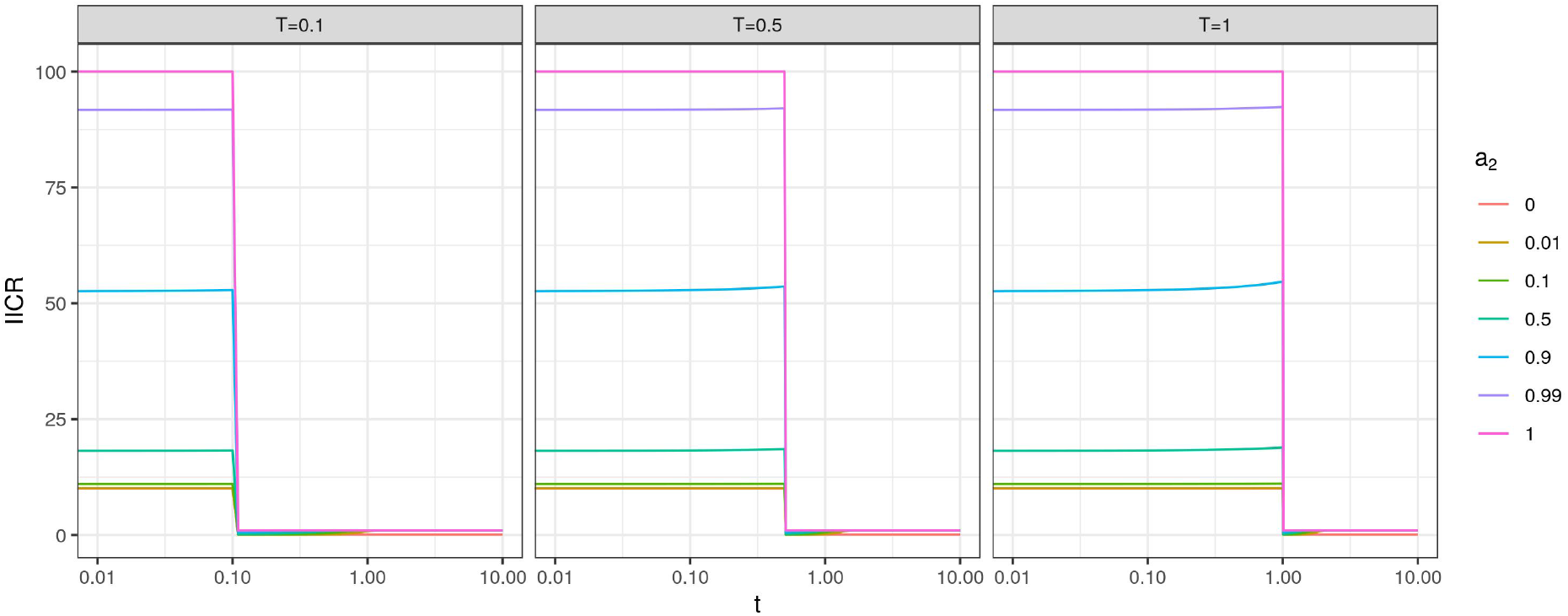
IICR curves for a panmictic model with a recent 100 fold expansion and *K* = 2 classes of genomic regions. Same as Figure 4 with a stronger population expansion (100 fold vs 5 fold).

**Figure S5:**
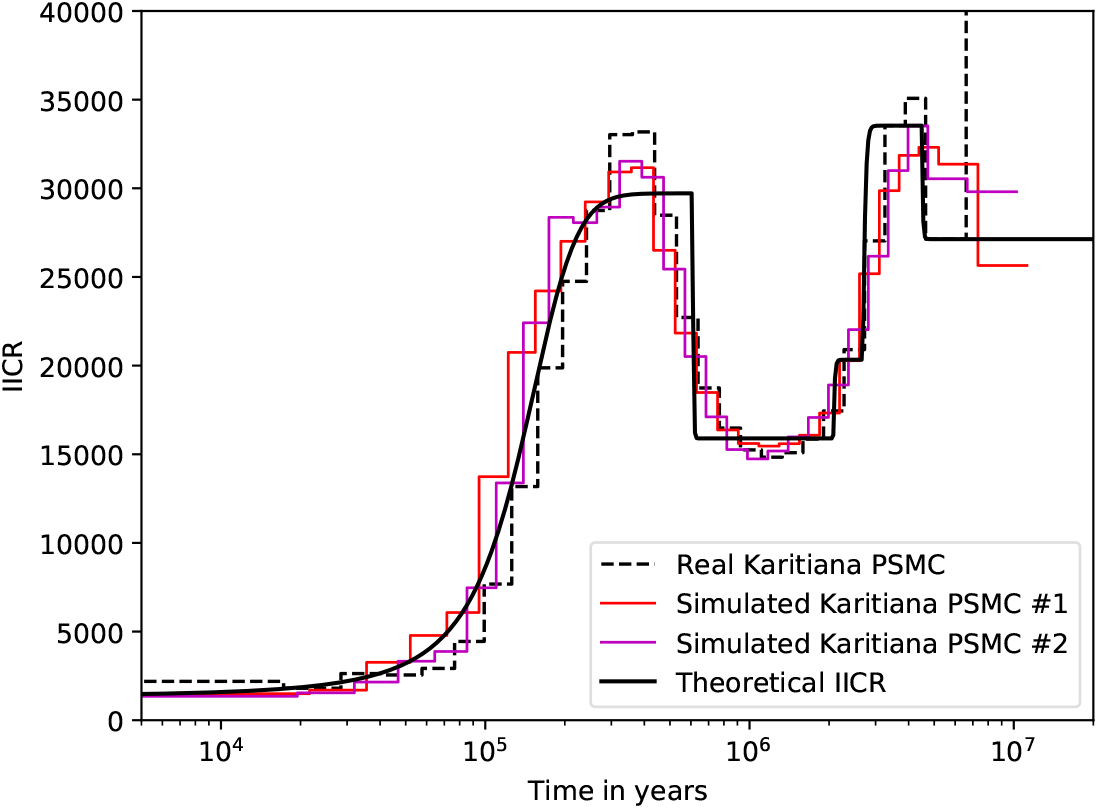
PSMC curves of simulated data under a non-stationary n-island model. We show in black the exact IICR corresponding to an inferred n-island model for a Karitiana individual in Arredondo et al. (2021). In color, we show various PSMC curves obtained by independently simulating genomic sequences under this structured model. The real PSMC curve for this Karitiana individual is represented by the dashed plot (Prado-Martinez et al., 2013). The horizontal axis is the time in years, with a generation time of 25 years. The vertical axis is the diploid population size. Times and population sizes were scaled assuming a mutation rate *μ*=1.25e-8.

**Figure S6:**
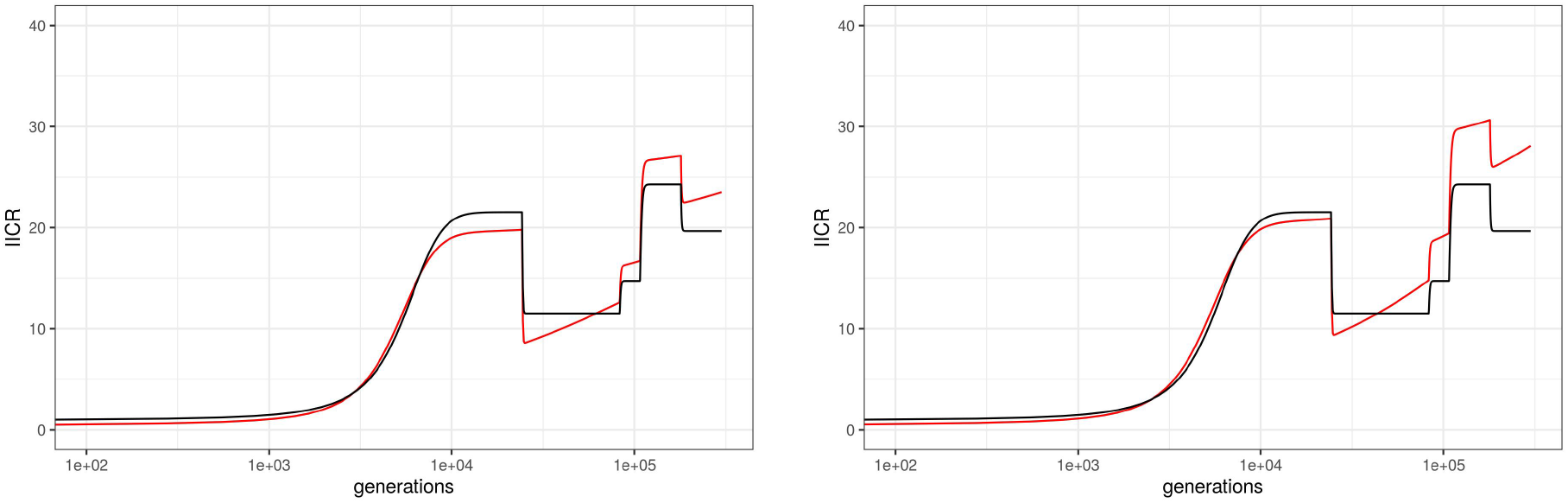
IICRs for demographic models combining population structure and linked selection in humans. Same as Figure 6, bottom panel, except that *λ* values greater than 2 (left) or 3 (right) were filtered out from the distribution in order to mimic a situation were loci under balancing selection could be detected and removed before computing the IICR. The resulting truncated distribution was rescaled.

